# A luciferase prosubstrate and a red bioluminescent calcium indicator to image neuronal activity

**DOI:** 10.1101/2021.12.06.471018

**Authors:** Xiaodong Tian, Yiyu Zhang, Xinyu Li, Ying Xiong, Tianchen Wu, Hui-wang Ai

## Abstract

Genetically encoded fluorescent indicators have been broadly used to monitor neuronal activity in live animals, but invasive surgical procedures are required. This study presents a functional bioluminescence imaging (fBLI) method for recording the activity of neuronal ensembles in the brain in awake mice. We developed a luciferase prosubstrate activatable *in vivo* by nonspecific esterase to enhance the brain delivery of the luciferin. We further engineered a bright, bioluminescent indicator with robust responsiveness to calcium ions (Ca^2+^) and appreciable emission above 600 nm. Integration of these advantageous components enabled the imaging of Ca^2+^ dynamics in awake mice minimally invasively with excellent signal-to-background and subsecond temporal resolution. This study thus establishes a new paradigm for studying brain functions in health and disease.

---

Fluorescence imaging is the standard method for neurobiologists to follow brain activity in small behaving animals. Genetically encoded fluorescent indicators, including GCaMPs and other Ca^2+^, voltage, neurotransmitter indicators, allowed the tracking of neuronal activities of specific brain regions and cell types in mammals with a high spatiotemporal resolution for extended periods.^1-3^ Despite the progress, fluorescence neuronal imaging is practically invasive and only reaches a shallow depth. Cranial windows or thinned skulls are often used to access the cortex.

Due to tissue absorption and scattering, the imaging depth is ∼ 200 μm for widefield one-photon and a couple of millimeters with multiphoton excitation.^3-5^ To reach deeper brain regions, more invasive procedures, such as implanting optical fibers or gradient-index (GRIN) lenses, are needed.^3^

Bioluminescence, which refers to photon emission from luciferase-catalyzed exothermic oxidation of the corresponding luciferin, is a promising imaging modality for noninvasive *in vivo* recording.^6,7^ Because bioluminescence needs no excitation, photons emitting from the embedded light sources can travel through several centimeters of mammalian tissue.^8^ In addition, compared to fluorescence, BLI has a low background, no photobleaching and phototoxicity, and minimized disturbances of light-sensitive biological components (*e*.*g*., the circadian system). Moreover, BLI has excellent compatibility with optogenetic and optochemical tools popular in neurobiology.

Commonly used luciferase-luciferin pairs originate from either insects or marine organisms.^6,7^ Insect luciferases are generally catalytically slow and consume adenosine triphosphate (ATP) for luciferin activation and photon production. In contrast, the oxidation of the coelenterazine (CTZ) luciferin by marine luciferases is ATP-independent. NanoLuc, a marine luciferase variant, exhibits a high photon production rate in the presence of furimazine (FRZ), a synthetic CTZ analog.^9^ However, NanoLuc has several unfavorable features for *in vivo* BLI, including low tissue penetration of its blue emission, and limited substrate solubility and stability. Recent studies have partially addressed some issues by developing new CTZ analogs and NanoLuc mutants^10-13^ or genetically fusing NanoLuc to long-wavelength-emitting fluorescent proteins (FPs) for redder emission via bioluminescence resonance energy transfer (BRET).^10,11,14,15^

Bioluminescent indicators that change signals in response to neuronal activity are needed for functional imaging. Ca^2+^ is a ubiquitous second messenger and intracellular Ca^2+^ has been used as a proxy for neuronal activity.^1-3^ Previous studies have used aequorin, a Ca^2+^-sensitive photoprotein, and its mutants for Ca^2+^ detection, but they all emit photons very slowly.^16,17^ Other studies introduced Ca^2+^-sensory elements into luciferases, including NanoLuc and NanoLuc-FP hybrid reporters, resulting in Ca^2+^ indicators with much-improved light production rates.^18-22^

Despite the progress, functional BLI (fBLI) of neuronal activity, which requires fast digitalization, is still hindered by insufficient photons reaching detectors. First, the intrinsic photon production rates of NanoLuc and NanoLuc-based indicators remain several orders of magnitude lower than those achievable in typical fluorescence imaging setups.^23,24^ Another limiting factor is the low amount of marine luciferase substrates that can be systematically delivered to the brain.^12,25,26^ Moreover, most current bioluminescent Ca^2+^ indicators (**Supplementary Table 1**) emit short-wavelength light strongly attenuated by brain tissue, skull, and skin.^19-22^ Orange CaMBIs, which were created by inserting the Ca^2+^-sensory calmodulin (CaM) and M13 moieties between residues 133 and 134 of NanoLuc linked to two copies of CyOFP1 (an orange-emitting FP), are the only NanoLuc-based indicators with appreciable emission above 600 nm.^24^ The Orange CaMBI 110 (OCaMBI110) variant has been used to image *in vivo* Ca^2+^ dynamics in the mouse liver but not yet in the brain.^12,24^

To fill the fBLI technical gap, we developed a luciferase prosubstrate for enhanced brain delivery and a bioluminescent indicator with redder emission and remarkable brightness and Ca^2+^ responsiveness. The integrated system drastically increased photon flux at the detector, enabling the minimally invasive imaging of deep-brain Ca^2+^ dynamics in awake mice responding to behavioral and disease triggers.

Our group previously reported teLuc, a bright and red-shifted NanoLuc mutant, and its paired DTZ substrate with promising BLI performance in mice.^10^ DTZ could be synthesized from inexpensive commercial reagents in two steps with excellent yields (**Supplementary Fig. 1**). First, we compared the computed logP (the octanol/water partitioning coefficient) of DTZ (∼4.3) with common BBB-permeable drugs.^27^ Decreasing the lipophilicity of DTZ was suggested to increase its delivery to the brain. Furthermore, although the mechanisms limiting the peripheral delivery of CTZ and its analogs to the brain are not fully understood, the P-glycoprotein (P-gp) efflux transporter on the blood-brain barrier (BBB) was shown to pump out CTZ.^28^ The BBB efflux issue is further compounded by other unfavorable factors, such as the rapid clearance and low solubility of these substrates.^25,29^ The literature reported that adding a succinate group to paclitaxel (Taxol) reduced the interaction with P-gp and increased brain distribution vastly,^30,31^ because P-gp unfavorably interacts with negatively charged molecules and the succinate addition installs a carboxylate functional group with a *p*K_a_ of ∼4. Thus, we chose to use the carboxylate functional group for modifying DTZ with the hope of enhancing hydrophilicity, reducing P-gp efflux, and increasing possible dosage via aqueous intravenous injection buffers.

Because the C3 carbonyl group of the DTZ imidazopyrazine ring (**Fig. 1A**) is required for substrate oxidation,^6^ C3 derivatizations will generate caged substrates resistant to auto- and luciferase-catalyzed oxidation. We designed and synthesized a compound with an extended carboxylate via C3 (see ETZ in **Fig. 1A** and **Supplementary Fig. 1**). When ETZ is delivered in vivo, nonspecific esterase is expected to hydrolyze the ester bonds, resulting in free DTZ for luciferase-catalyzed bioluminescence (**Fig. 1B**). We named this new compound ETZ for <u>e</u>sterase-dependent activation and <u>e</u>nhanced *in vivo* performance (presented below). For comparison purposes, we synthesized C3-DMA-DTZ (**Fig. 1A** and **Supplementary Fig. 1**), which contains ester linkages but is positively charged at physiological *p*H.

**Fig. 1.**
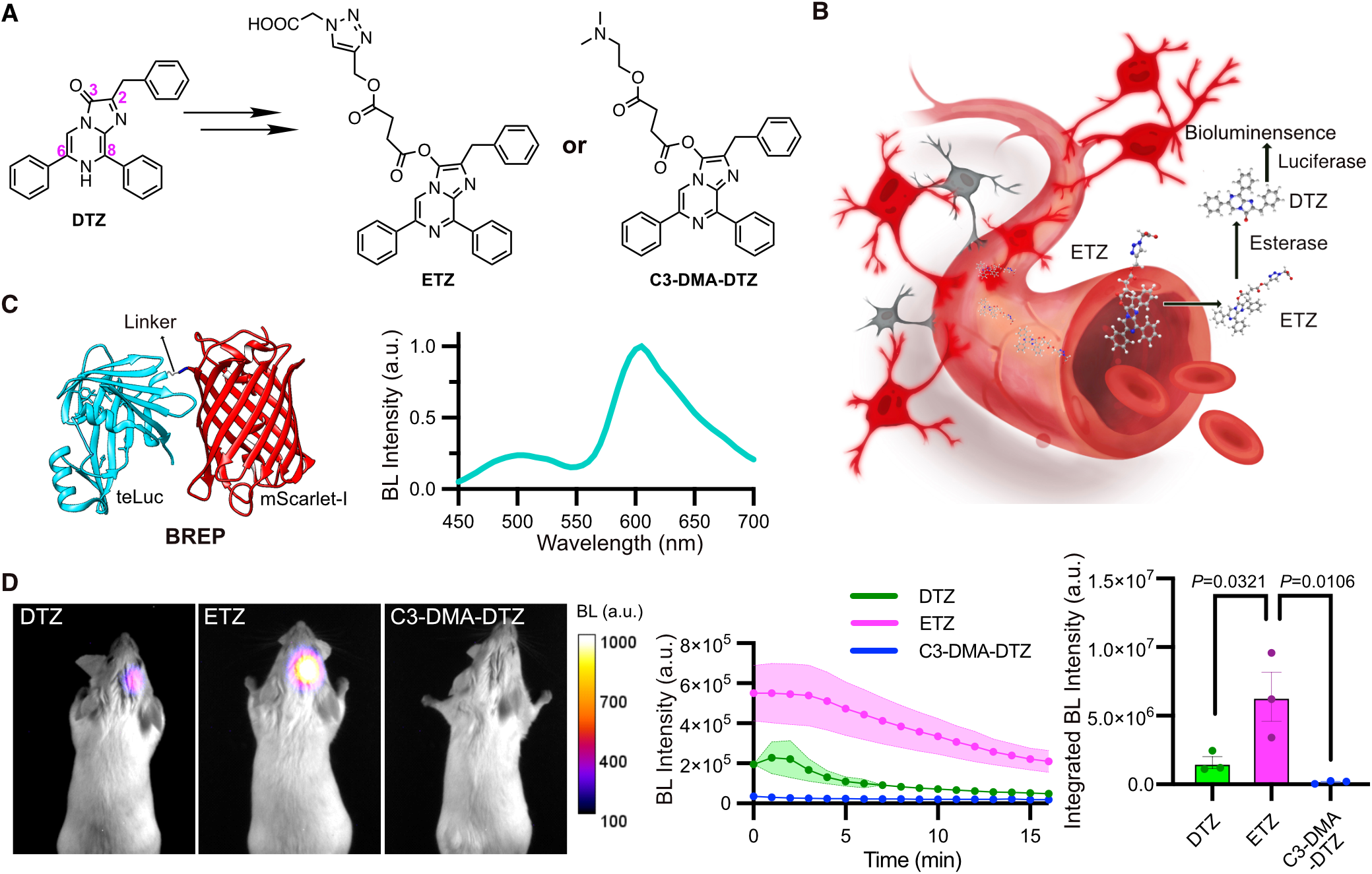
New luciferase prosubstrates and initial evaluation for brain imaging with BREP. (**A**) Illustration of the derivatization of DTZ to generate two luciferase prosubstrates (ETZ and C3-DMA-DTZ) with modulated solubility, pharmacokinetics, and brain permeability and retention. (**B**) Proposed conversion of ETZ to DTZ *in vivo* by nonspecific esterase, followed by luciferase-catalyzed oxidation of DTZ to generate bioluminescence. (**C**) Schematic illustration of the domain arrangement of BREP (left) and its bioluminescence emission spectrum in the presence of DTZ (right). (**D**) Left: Representative bioluminescence images of live mice with BREP-expressing HEK 293T cells stereotactically injected into the hippocampus. The substrates were administered via tail vein at their respective saturation concentrations. Images with peak bioluminescence intensities were presented in pseudocolor overlaid on corresponding brightfield images. Middle: Bioluminescence intensity over time shown for each substrate. Right: Comparison of the integrated bioluminescence intensity (area under the curve) with the residual background subtracted. Data are presented as mean ± sem (n=3 mice). *P* values were derived from ordinary one-way ANOVA followed by Dunnett’s multiple comparisons test.

We previously fused a LumiLuc luciferase to a bright red FP (RFP) mScarlet-I, resulting in a LumiScarlet reporter with ∼ 51% of the total emission above 600 nm.^11^ Following the success, we created a similar fusion between teLuc and mScarlet-I and optimized the linker for increased BRET efficiency (**Supplementary Fig. 2**). We arrived at a bright <u>B</u>ioluminescent <u>Re</u>d <u>P</u>rotein (BREP) with ∼60% of its total emission above 600 nm (**Fig. 1C**). Although teLuc is less red-shifted than LumiLuc and has less spectral overlap with mScarlet-I, we observed more effective BRET in BREP than LumiScarlet due to the shorter donor-acceptor distance and a possible spatial orientation favoring donor-acceptor dipole coupling in BREP.^32^

We next compared DTZ, ETZ, and C3-DMA-DTZ for brain delivery in mice. We stereotactically injected 7,000 human embryonic kidney (HEK) 293T cells transiently transfected with BREP to the hippocampus in anesthetized BALB/cJ mice. We immediately administered 100 μL of buffers containing each substrate at their saturation concentrations via tail vein. ETZ exhibited better solubility than DTZ and C3-DMA-DTZ (**Supplementary Fig. 3A**) and could be delivered at a dosage of 0.68 µmol per mouse. Mice infused with ETZ showed the most robust and durable bioluminescence (**Fig. 1D** and **Supplementary Fig. 3B**). During the 16-min window used in this experiment, the integrated signal of the ETZ group was ∼4.0-fold and ∼39.0-fold higher than the DTZ and C3-DMA-DTZ groups, respectively. The results suggest that ETZ is a promising substrate for brain imaging in live mice.

We further engineered a Ca^2+^ indicator from BREP. Inspired by Orange CaMBIs, we similarly inserted CaM and M13 between residues 133 and 134 of teLuc in BREP, resulting in a prototype showing 2.5-fold (BL/BL_0_) Ca^2+^-dependent bioluminescence increase (**Supplementary Fig. 4**). Next, we performed six rounds of error-prone PCRs and screened the libraries for high bioluminescence brightness and Ca^2+^ responsiveness. The effort led to a <u>B</u>ioluminescent <u>R</u>ed <u>I</u>ndicator for <u>C</u>a^2+^ (BRIC) with a 6.5-fold (BL/BL_0_) response (**Fig. 2AB, Supplementary Fig. 5**, and **Supplementary Table 1**). BRIC, at its Ca^2+^-bound condition, retained ∼ 46% of the brightness of BREP. Using the purified protein, the dissociation constant (*K*_d_) of BRIC to Ca^2+^ was determined to be 133 nM (**Fig. 2C**). The response magnitude increased as pH changed from 5.5 to 7 and was relatively stable at pH > 7 (**Fig. 2D**). When compared in parallel, BRIC was more responsive to Ca^2+^ than OCaMBI110 (**Supplementary Fig. 6AB** and **Supplementary Table 1)** To evaluate BRIC for imaging cellular Ca^2+^ dynamics, we transiently expressed BRIC in human cervical cancer HeLa cells, in which histamine can evoke Ca^2+^ waves.^33^ As expected, we observed single-cell bioluminescence oscillations in response to histamine under bioluminescence microscopy (**Fig. 2E** and **Supplementary Movie 1**). Furthermore, we prepared adeno-associated viruses (AAVs) with BRIC expression under the human synapsin I (hSyn) promoter. We transduced cultured primary mouse neurons and successfully detected Ca^2+^ influx after high K^+^ depolarization, while the control BREP-expressing neurons showed little bioluminescence increase (**Fig. 2F** and **Supplementary Movie 2**).

**Fig. 2.**
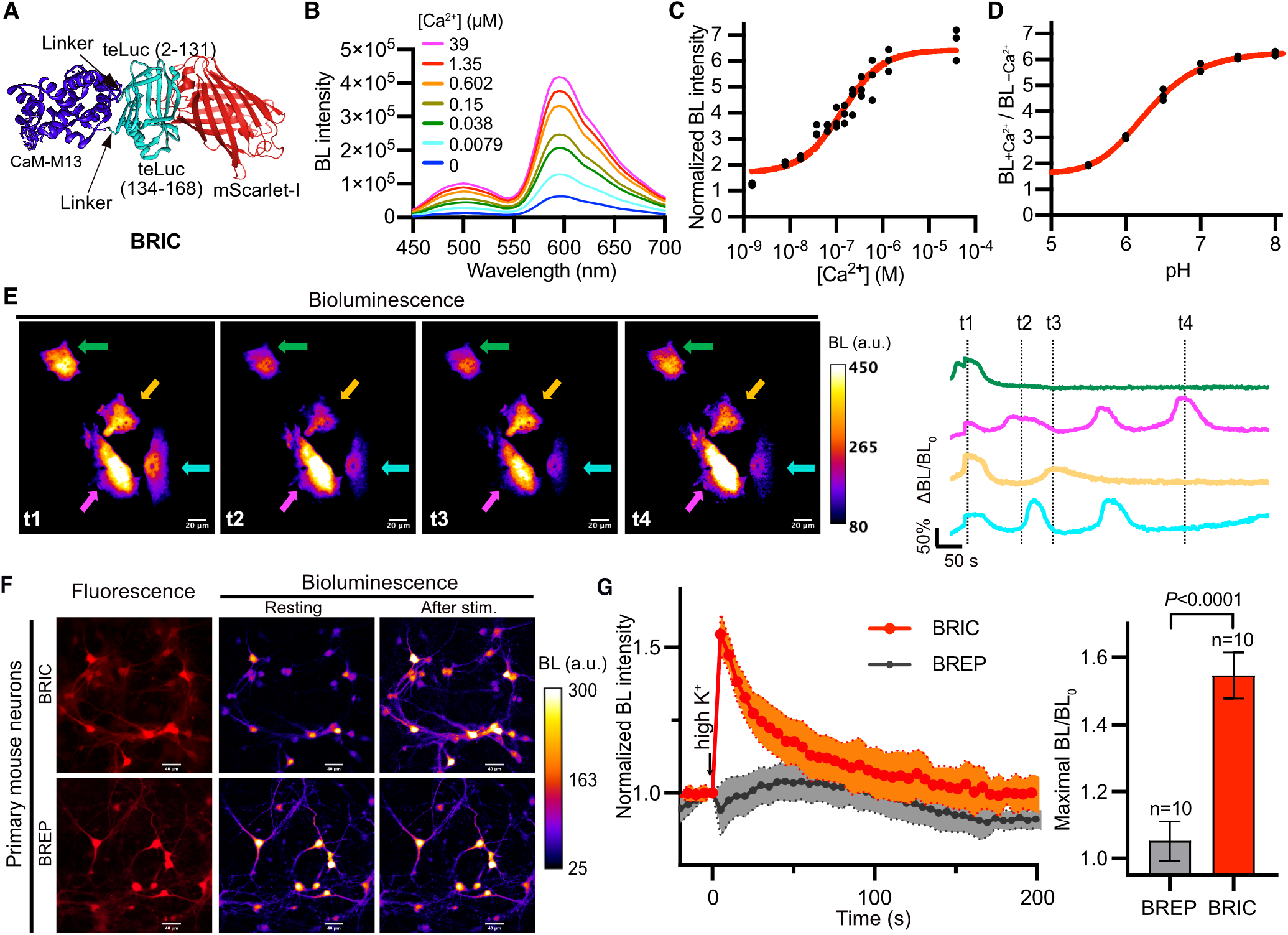
Characterization of BRIC in vitro and cultured cells. (**A**) Schematic illustration of the domain arrangement of BRIC. (**B**) Bioluminescence spectra of BRIC in the presence of DTZ and indicated concentrations of free Ca^2+^. (**C**) Ca^2+^-dependency of BRIC bioluminescence at the peak emission wavelength (595 nm). n=3. A one-site binding model was used to fit the data and derive the dissociation constant (*K*_d_ = 133±24 nM). (**D**) Ratios of BRIC bioluminescence in the presence (39 μM) to the absence of Ca^2+^ across the indicated pH range. n=3. (**E**) Representative pseudocolored bioluminescence images (left) and intensity traces (right) of histamine-induced Ca^2+^ dynamics in HeLa cells. Arrows indicate individual cells, and the colors of the arrows are identical to the colors of the intensity traces. The baselines of the intensity traces were corrected for monoexponential decay caused by substrate consumption. The experiment was repeated five times with similar results. Scale bar, 20 μm. (**F**) Representative fluorescence and bioluminescence images of BRIC- or BREP-expressing primary mouse neurons. High K^+^ (30 mM) was used to depolarize the neurons. Scale bar, 40 μm. (**G**) Quantification of bioluminescence intensity changes of neurons in response to high K^+^. The baselines were corrected using a monoexponential decay model. Data are presented as mean ± s.d., and the *P* value was derived from unpaired two-tailed *t*-tests.

We compared BRIC with OCaMBI110 in HeLa cells and cultured mouse neurons. Under fluorescence channels, we observed extensive puncta in cells overexpressing OCaMBI110 (**Supplementary Fig. 6C**) but not in BRIC-expressing cells. In addition, the cells with fluorescent puncta showed little bioluminescence and were unresponsive to histamine or high K^+^. Although the exact reason is unknown, the observed puncta may be OCaMBI110 oligomers because each OCaMBI110 molecule contains two copies of dimeric CyOFP1 (**Supplementary Fig. 6D**).^14^ Moreover, with punctum-containing cells excluded from analysis, the response magnitude of BRIC was still higher than OCaMBI110 in both HeLa cells and cultured neurons (**Supplementary Fig. 6EF**).

We stereotactically injected BRIC and OCaMBI110 AAVs (adjusted to the same viral titer) into the hippocampus of BALB/cJ mice (**Fig. 3A**) and compared the brightness of the two indicators in day 19 post viral administration. FFz was recently reported as a new NanoLuc substrate with enhanced *in vivo* performance,^12^ so we chemically synthesized FFz and examined OCaMBI110 with either FRZ or FFz. The substrate solubility in the injection buffers was determined (**Supplementary Fig. 7A**), and each substrate was intravenously administered to anesthetized animals at their saturation concentrations. Using an EMCCD camera, we followed the signals of BRIC with 500-ms exposure for 1 h. The starting bioluminescence of BRIC in the presence of ETZ was ∼ 168- and 29-fold higher than OCaMBI110 in the presence of FRZ and FFz, respectively (**Fig. 3B** and **Supplementary Fig. 7B**). The BRIC signals were consistently higher than the background during periods much longer than OCaMBI110 with either substrate (**Fig. 3C** and **Supplementary Fig. 7C**). In terms of the signals integrated over time, BRIC was ∼ 153.7- and 22.0-fold of OCaMBI110 in the presence of FRZ and FFz, respectively (**Fig. 3D**). Next, we prepared acute brain slices from BRIC-expressing mice and observed bioluminescence rise in response to high K^+^, confirming the activity of BRIC in the brain tissue (**Supplementary Fig. 8** and **Supplementary Movie 3**).

**Fig. 3.**
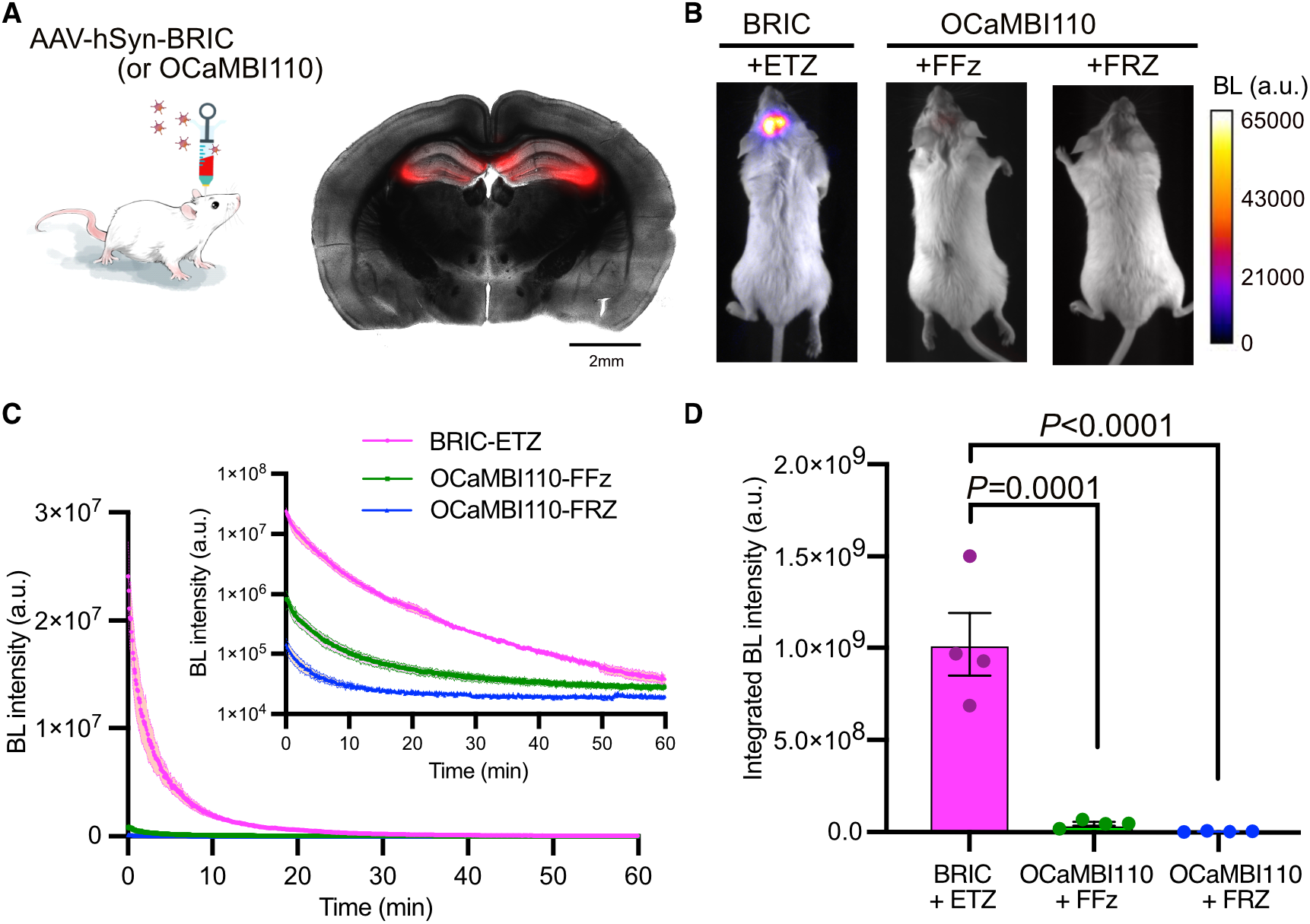
Brightness comparison of BRIC and Orange CaMBI 110 (OCaMBI110) in the hippocampus in live mice. (**A**) Left: Illustration of stereotactic intracranial administration of AAVs. Right: Image of an acute brain slice prepared from a BRIC-transduced mouse, showing the successful expression of the indicator in the hippocampus. The fluorescence channel (red) is overlaid on the corresponding grayscale brightfield image. Scale bar, 2 mm. (**B**) Representative bioluminescence images of live mice with the hippocampus transduced with BRIC or OCaMBI110 AAVs. The substrates were administered via tail vein at their respective saturation concentrations. Images with peak bioluminescence intensities were presented in pseudocolor overlaid on corresponding brightfield images. (**C**) Bioluminescence intensity over time presented for each substrate. The inset shows the same results with a logarithmic y-axis. (**D**) Comparison of the integrated bioluminescence intensity (area under the curve) with the residual background subtracted. Data are presented as mean ± sem (n=4 mice). *P* values were derived from ordinary one-way ANOVA followed by Dunnett’s multiple comparisons test.

We finally examined BRIC for monitoring Ca^2+^ dynamics in awake mice (**Fig. 4A**). First, we administered the virus to the basolateral amygdala (BLA) region (**Supplementary Fig. 9**) responsible for fear processing.^34^ Upon the intravenous injection of ETZ, head-fixed mice were subjected to BLI. In response to footshock stimuli, we observed reproducible bioluminescence increases from BRIC-expressing mice (**Fig. 4B** and **Supplementary Movie 4**). In addition, the indicator was expressed in the hippocampus of C57BL/6J mice, and kainic acid (KA) was used to induce epileptic seizures, a condition known for abnormal hippocampal Ca^2+^ waves.^35,36^ As expected, we detected rapid bioluminescence intensity changes from BRIC-expressing mice (**Fig. 4C** and **Supplementary Movie 5**). BREP was included as a negative control, and BREP-expressing mice showed little response to footshock and during seizures.

**Fig. 4.**
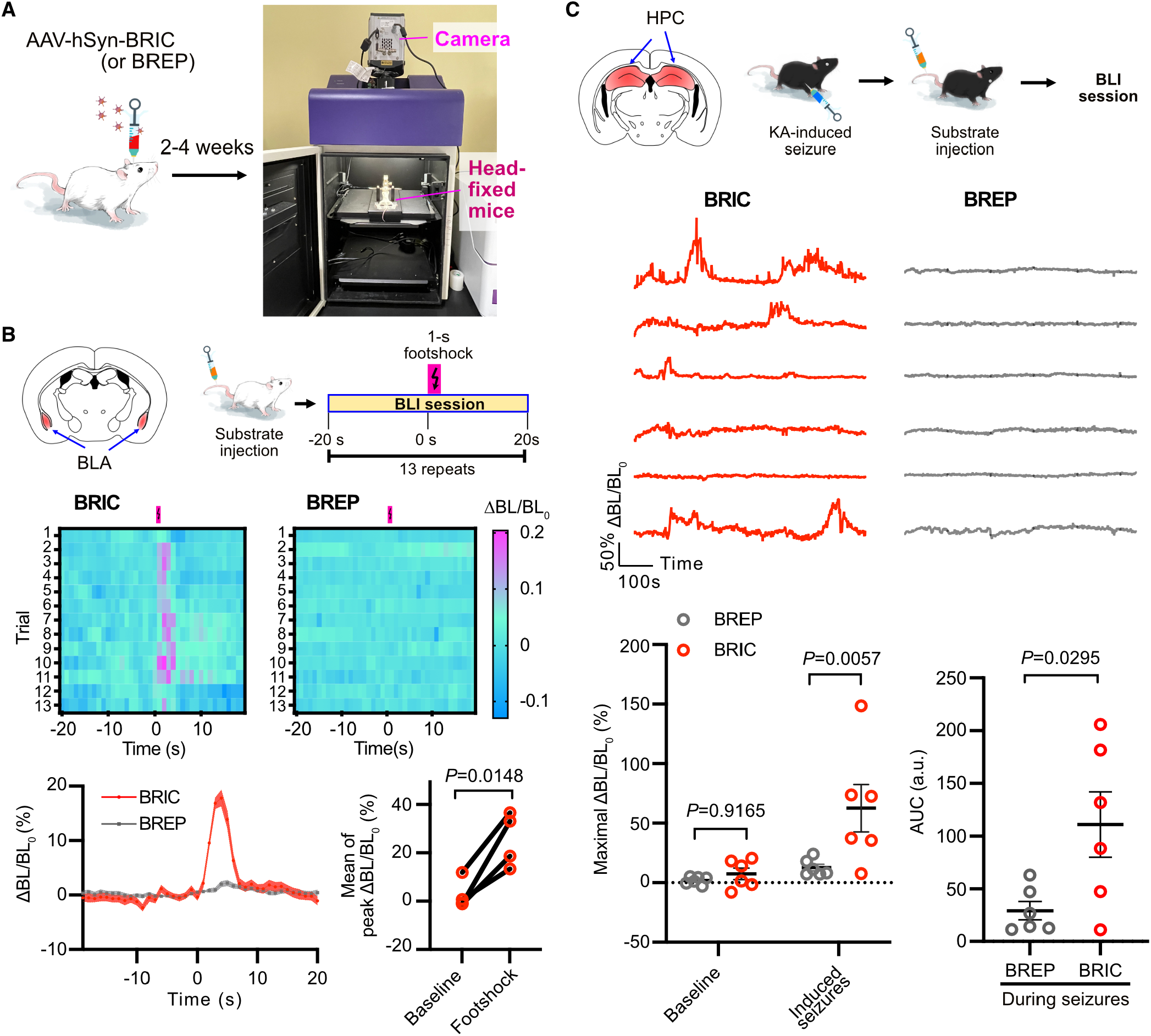
Bioluminescence imaging of Ca^2+^ dynamics in the brain in awake mice. (**A**) Illustration of intracranial viral administration and bioluminescence imaging of head-fixed awake mice. (**B**) Bioluminescence imaging of footshock-induced Ca^2+^ in the basolateral amygdala (BLA). Top: Viral injection sites and the general experimental procedure. Middle: Bioluminescence intensity heatmap of a representative BRIC- or BREP-expressing mouse in response to 13 consecutive trials of footshock stimulations. Bottom: Quantification of bioluminescence intensity changes. Data are presented as mean ± sem (n=4 mice, each with 13 trials). The right panel compares the average responses of 13 trials for each mouse. The *P* value was derived from paired two-tailed *t*-tests. (**C**) Bioluminescence imaging of Ca^2+^ in the hippocampus (HPC) during kainic acid (KA)-induced seizures. Top: Viral injection sites and the general experimental procedure. Middle: Bioluminescence intensity traces of BRIC- or BREP-expressing mouse during seizures. Bottom: Comparison of maximal intensity changes and area under the curve (AUC). Data are presented as mean ± sem. n=6 mice. *P* values were derived from two-way ANOVA followed by Šídák’s multiple comparisons test or unpaired two-tailed *t*-tests

In summary, we have developed an integrated fBLI platform for imaging brain Ca^2+^ dynamics in awake mice. We first enhanced the brain delivery of the luciferase substrate via a prosubstrate strategy. We next developed BREP, a luciferase-FP fusion reporter, with remarkable emission above 600 nm. From BREP, we further engineered a bioluminescent Ca^2+^ indicator with high brightness and Ca^2+^ responsiveness. Finally, we validated the resultant system in cultured cells, brain slices, and live animals. BLI only requires a dark box with a sensitive camera. Similar setups are already in many laboratories and core facilities; otherwise, they are commercially available or can be constructed at affordable costs. Although the new paradigm cannot provide spatial information as detailed as fluorescence neuronal imaging, this less invasive method would be suitable for studying the functions of neuronal populations in behaving animals.

CTZ has been used to activate luminopsins (luciferase-channelrhodopsin fusions) in the brain in live mice,^37^ but a fast clearance of CTZ from the brain is advantageous in that case since it facilitates temporal control. In this study, we introduced a carboxylate via the C3 position of the DTZ imidazopyrazine ring, leading to the ETZ prosubstrate exhibiting resistance to auto-oxidation, increased aqueous solubility, and extended *in vivo* kinetics. When administered to mice, the labile ester linkages of ETZ may break down before it crosses the BBB. We thus do not know whether or how much the P-gp efflux pump plays a role in enhancing the brain delivery of the substrate, although it was an initial inspiration for our substrate designing. Future studies may modify luciferins with a stable carboxylate functional group, but it will require further engineering of corresponding luciferases and indicators to accommodate the new substrates.

We had to apply corrections for baseline intensity decays caused by the substrate loss for time-lapse quantitation. A monoexponential decay model was adequate for baseline corrections in those experiments spanning short periods, although a more complex decay was evident during the more extended 60-min period shown in **Fig. 3C**. Future studies may allow BLI with two channels above 600 nm, enabling ratiometric correction for baseline decays and motions of freely moving animals.

In addition, a recent study described the use of FRZ and a NanoLuc-derived bioluminescent voltage indicator for imaging cortical activity in mice.^26^ To gain enough signals, a cranial window was used for bioluminescence collection and continuous substrate delivery to the cortex. Furthermore, a GaAsP image intensifier was placed in front of the EMCCD to boost signals.

Another recent preprint described a NanoLuc-derived bioluminescent glutamate indicator that has been tested in cultured cells.^38^ We expect our results and strategies presented here to assist in the further development of these bioluminescent indicators for deep-brain *in vivo* imaging with minimally invasive procedures. Furthermore, as much enhanced BLI tools, ETZ, BREP, and BRIC are expected to find applications and generate impacts beyond neurobiology.

## Acknowledgments

We thank Dr. Zefan Li for confirming the result presented in Fig. 2B, and other members in the Ai lab for helpful discussion and assistance with experiments. We also acknowledge the UVA NMR Spectroscopy Core Facility for technical assistance. Research reported in this publication was supported by the University of Virginia Start-up Fund and National Institutes of Health grants (R01DK122253 and R01GM129291) to HA.

## Author contributions

HA conceived the project. XT synthesized compounds, engineered and characterized BRIC *in vitro* and in cultured cells, and prepared virus. YZ performed *in vivo* viral injection. XL prepared primary neurons. XT, XL and YZ performed *in vivo* brightness comparison of substrates and indicators. YZ and XT imaged brain slices and live animals. TW prepared some brain slices and conducted intravenous injections of substrates. YX developed and recorded the emission of BREP. XT, YZ, XL and YX analyzed data and prepared figures. HA, XT, YZ, and YX wrote the manuscript.

## References

1 Sabatini, B. L. & Tian, L. Imaging Neurotransmitter and Neuromodulator Dynamics In Vivo with Genetically Encoded Indicators. Neuron 108, 17–32, doi:10.1016/j.neuron.2020.09.036 (2020).

2 Chen, T. W. et al. Ultrasensitive fluorescent proteins for imaging neuronal activity. Nature 499, 295–300, doi:10.1038/nature12354 (2013).

3 Campos, P., Walker, J. J. & Mollard, P. Diving into the brain: deep-brain imaging techniques in conscious animals. J. Endocrinol. 246, R33–R50, doi:10.1530/joe-20-0028 (2020).

4 Miller, D. R., Jarrett, J. W., Hassan, A. M. & Dunn, A. K. Deep Tissue Imaging with Multiphoton Fluorescence Microscopy. Curr. Opin. Biomed. Eng. 4, 32–39, doi:10.1016/j.cobme.2017.09.004 (2017).

5 Aharoni, D. & Hoogland, T. M. Circuit Investigations With Open-Source Miniaturized Microscopes: Past, Present and Future. Front. Cell. Neurosci. 13, 141, doi:10.3389/fncel.2019.00141 (2019).

6 Yeh, H. W. & Ai, H. W. Development and Applications of Bioluminescent and Chemiluminescent Reporters and Biosensors. Annu. Rev. Anal. Chem. 12, 129–150, doi:10.1146/annurev-anchem-061318-115027 (2019).

7 Yao, Z., Zhang, B. S. & Prescher, J. A. Advances in bioluminescence imaging: new probes from old recipes. Curr. Opin. Chem. Biol. 45, 148–156, doi:10.1016/j.cbpa.2018.05.009 (2018).

8 Sadikot, R. T. & Blackwell, T. S. Bioluminescence imaging. Proc. Am. Thorac. Soc. 2, 537–540, doi:10.1513/pats.200507-067DS (2005).

9 Hall, M. P. et al. Engineered luciferase reporter from a deep sea shrimp utilizing a novel imidazopyrazinone substrate. ACS Chem. Biol. 7, 1848–1857, doi:10.1021/cb3002478 (2012).

10 Yeh, H. W. et al. Red-shifted luciferase-luciferin pairs for enhanced bioluminescence imaging. Nat. Methods 14, 971–974, doi:10.1038/nmeth.4400 (2017).

11 Yeh, H. W. et al. ATP-Independent Bioluminescent Reporter Variants To Improve in Vivo Imaging. ACS Chem. Biol. 14, 959–965, doi:10.1021/acschembio.9b00150 (2019).

12 Su, Y. et al. Novel NanoLuc substrates enable bright two-population bioluminescence imaging in animals. Nat. Methods 17, 852–860, doi:10.1038/s41592-020-0889-6 (2020).

13 Shakhmin, A. et al. Coelenterazine analogues emit red-shifted bioluminescence with NanoLuc. Org. Biomol. Chem. 15, 8559–8567, doi:10.1039/c7ob01985h (2017).

14 Chu, J. et al. A bright cyan-excitable orange fluorescent protein facilitates dual-emission microscopy and enhances bioluminescence imaging in vivo. Nat. Biotechnol. 34, 760–767, doi:10.1038/nbt.3550 (2016).

15 Schaub, F. X. et al. Fluorophore-NanoLuc BRET Reporters Enable Sensitive In Vivo Optical Imaging and Flow Cytometry for Monitoring Tumorigenesis. Cancer Res. 75, 5023–5033, doi:10.1158/0008-5472.Can-14-3538 (2015).

16 Prasher, D., McCann, R. O. & Cormier, M. J. Cloning and expression of the cDNA coding for aequorin, a bioluminescent calcium-binding protein. Biochem. Biophys. Res. Commun. 126, 1259–1268 (1985).

17 Curie, T., Rogers, K. L., Colasante, C. & Brûlet, P. Red-shifted aequorin-based bioluminescent reporters for in vivo imaging of Ca^2+^ signaling. Mol. Imaging 6, 30–42 (2007).

18 Saito, K. et al. Luminescent proteins for high-speed single-cell and whole-body imaging. Nat. Commun. 3, 1262, doi:10.1038/ncomms2248 (2012).

19 Suzuki, K. et al. Five colour variants of bright luminescent protein for real-time multicolour bioimaging. Nat. Commun. 7, 13718, doi:10.1038/ncomms13718 (2016).

20 Yang, J. et al. Coupling optogenetic stimulation with NanoLuc-based luminescence (BRET) Ca^2+^ sensing. Nat. Commun. 7, 13268, doi:10.1038/ncomms13268 (2016).

21 Qian, Y., Rancic, V., Wu, J., Ballanyi, K. & Campbell, R. E. A Bioluminescent Ca^2+^ Indicator Based on a Topological Variant of GCaMP6s. ChemBioChem 20, 516–520, doi:10.1002/cbic.201800255 (2019).

22 Farhana, I., Hossain, M. N., Suzuki, K., Matsuda, T. & Nagai, T. Genetically Encoded Fluorescence/Bioluminescence Bimodal Indicators for Ca^2+^ Imaging. ACS Sens. 4, 1825–1834, doi:10.1021/acssensors.9b00531 (2019).

23 Hess, S. T., Girirajan, T. P. & Mason, M. D. Ultra-high resolution imaging by fluorescence photoactivation localization microscopy. Biophys. J. 91, 4258–4272, doi:10.1529/biophysj.106.091116 (2006).

24 Oh, Y. et al. An orange calcium-modulated bioluminescent indicator for non-invasive activity imaging. Nat. Chem. Biol. 15, 433–436, doi:10.1038/s41589-019-0256-z (2019).

25 Otto-Duessel, M. et al. In vivo testing of Renilla luciferase substrate analogs in an orthotopic murine model of human glioblastoma. Mol. Imaging 5, 57–64 (2006).

26 Inagaki, S. et al. Imaging local brain activity of multiple freely moving mice sharing the same environment. Sci. Rep. 9, 7460, doi:10.1038/s41598-019-43897-x (2019).

27 Pajouhesh, H. & Lenz, G. R. Medicinal chemical properties of successful central nervous system drugs. NeuroRx 2, 541–553, doi:10.1602/neurorx.2.4.541 (2005).

28 Pichler, A., Prior, J. L. & Piwnica-Worms, D. Imaging reversal of multidrug resistance in living mice with bioluminescence: MDR1 P-glycoprotein transports coelenterazine. Proc. Natl. Acad. Sci. USA 101, 1702–1707, doi:10.1073/pnas.0304326101 (2004).

29 Taylor, A., Sharkey, J., Plagge, A., Wilm, B. & Murray, P. Multicolour In Vivo Bioluminescence Imaging Using a NanoLuc-Based BRET Reporter in Combination with Firefly Luciferase. Contrast Media Mol. Imaging 2018, 2514796, doi:10.1155/2018/2514796 (2018).

30 Turunen, B. J. et al. Paclitaxel succinate analogs: Anionic and amide introduction as a strategy to impart blood-brain barrier permeability. Bioorg. Med. Chem. Lett. 18, 5971–5974, doi:10.1016/j.bmcl.2008.09.103 (2008).

31 Rice, A. et al. Chemical modification of paclitaxel (Taxol) reduces P-glycoprotein interactions and increases permeation across the blood-brain barrier in vitro and in situ. J. Med. Chem. 48, 832–838, doi:10.1021/jm040114b (2005).

32 Day, R. N. & Davidson, M. W. Fluorescent proteins for FRET microscopy: monitoring protein interactions in living cells. Bioessays 34, 341–350, doi:10.1002/bies.201100098 (2012).

33 Sauvé, R. et al. Ca^2+^ oscillations induced by histamine H1 receptor stimulation in HeLa cells: Fura-2 and patch clamp analysis. Cell Calcium 12, 165–176, doi:10.1016/0143-4160(91)90018-a (1991).

34 Sun, Y., Gooch, H. & Sah, P. Fear conditioning and the basolateral amygdala. F1000Res 9, doi:10.12688/f1000research.21201.1 (2020).

35 Berdyyeva, T. K. et al. Direct Imaging of Hippocampal Epileptiform Calcium Motifs Following Kainic Acid Administration in Freely Behaving Mice. Front. Neurosci. 10, 53, doi:10.3389/fnins.2016.00053 (2016).

36 Zhang, X. et al. Stereotypical patterns of epileptiform calcium signal in hippocampal CA1, CA3, dentate gyrus and entorhinal cortex in freely moving mice. Sci. Rep. 9, 4518, doi:10.1038/s41598-019-41241-x (2019).

37 Berglund, K. et al. Luminopsins integrate opto- and chemogenetics by using physical and biological light sources for opsin activation. Proc. Natl. Acad. Sci. U. S. A. 113, E358–367, doi:10.1073/pnas.1510899113 (2016).

38 Petersen, E. D. et al. Bioluminescent Genetically Encoded Glutamate Indicator for Molecular Imaging of Neuronal Activity. bioRxiv, DOI: 10.1101/2021.1106.1116.448690, doi:10.1101/2021.06.16.448690 (2021).

