## Supporting Information for "A luciferase prosubstrate and a red bioluminescent calcium indicator to image neuronal activity"

#### METHODS

**Reagents and General Methods.** Unless otherwise stated, all chemicals were purchased from Sigma-Aldrich, Fisher Scientific, or VWR and used without further purification. Kainic acid monohydrate (KA) was purchased from Cayman Chemical. pcDNA3.1-Orange_CaMBI_110 (Addgene #124094) and pAAV.hSyn.iGluSnFr.WPRE.SV40 (Addgene #98929) were gifts from M.Z. Lin and L.L. Looger, respectively. pAdDeltaF6 (Addgene #112867), pAAV2/9n (Addgene #112865) were gifts from J.M. Wilson. Synthetic DNA oligonucleotides were purchased from Integrated DNA Technologies or Eurofins Genomics. Restriction endonucleases and Phusion High-Fidelity DNA Polymerase were purchased from Thermo Scientific. Products of PCR and restriction digestion were purified with preparative agarose gel electrophoresis, followed by gel extraction. DNA sequences were analyzed by Eurofins Genomics. Gibson assembly was performed using a homemade kit by following a procedure from Addgene. Small-scale plasmid DNA preparation was performed using Miniprep kits from Syd Labs. Large-scale plasmid DNA preparation was performed using alkaline lysis followed by isopropanol precipitation, PEG 8000 precipitation, and phenol/chloroform extraction. Merck Geduran Si 60 silica gel was used for normal-phase column chromatography. A Waters Prep 150/SQ Detector 2 LC-MS Purification System equipped with an XBridge BEH Amide/Phenyl OBD Prep Column (130Å, 5 µm, 30 mm X 150 mm) was used for preparative reverse-phase HPLC purification. Lyophilization was performed on a 12-port Labconco freeze dryer with an Edwards RV3 vacuum pump. All NMR spectra were collected using a Bruker Avance III 600 MHz NMR spectrometer. Reference values for residual solvents were taken as 7.27 ($CDCl_3$) or 3.30 (methanol-$d_4$) ppm for $^1$H-NMR, and 77.2 ($CDCl_3$) or 49.0 (methanol-$d_4$) ppm for $^{13}$C-NMR. Splitting patterns of NMR active peaks are reported as s (singlet), d (doublet), t (triplet), dd (doublet of doublets), dt (doublet of triplets), and m (multiplet). All animal experiments were conducted according to the approval and guidelines of the University of Virginia Institutional Animal Care and Use Committee.

**Chemical Synthesis.** Synthetic schemes and compound numbering information are shown in **Supplementary Fig. 1**. The detailed procedures for compound synthesis are described below. Furimazine (FRZ) and FFz were synthesized following reported methods.[9,12]

**1,1-diethoxy-3-phenylacetone (Compound 1):** Compound **1** was synthesized by modifying a previous procedure.[39] Ethyl diethoxyacetate (10 mL, 56.8 mmol, 1.0 equiv.) was added into 30 mL anhydrous THF under argon (Ar) protection. Subsequently, BnMgCl (45 mL, 1.4 M in THF, 1.1 equiv.) was dropwise added into the reaction mixture in a dry ice-acetone bath. The mixture was stirred for 1 h in the dry ice-acetone bath, and then at room temperature for another 2 h. The reaction was next quenched by addition of $H_2O$. The mixture was filtrated and extracted three times with EtOAc. The organic layer was combined, dried with anhydrous $Na_2SO_4$, concentrated *in vacuo*. Purification was performed with silica gel chromatography (EtOAc/hexane = 1/5, v/v) to give compound **1** (8.8 g, 70.0 %).

**4-Oxo-4-(prop-2-yn-1-yloxy) butanoic acid (Compound 2):** NHS (920.0 mg, 8.0 mmol, 0.4 equiv.), DMAP (260.0 mg, 2.0 mmol, 0.1 equiv.), TEA (840.0 µL, 6.0 mmol, 0.3 equiv.) and propargyl alcohol (7.4 mL, 60.0 mmol, 3.0 equiv.) were separately added into a solution of succinic anhydride (2000.0 mg, 20.0 mmol, 1.0 equiv.) dissolved in 50 mL anhydrous toluene. The mixture was heated to reflux overnight under Ar. After cooling, the mixture was washed with brine and extracted three times with EtOAc. The organic layer was combined and dried with anhydrous $Na_2SO_4$. After filtration and rotovap concentration, the residue was further purified with silica gel chromatography (EtOAc/hexane/acetic acid = 1/4/0.005, v/v/v) to afford compound **2** (2028.0 mg, 65%).

**2-Amino-3,5-diphenylpyrazine (Compound 3):** Compound **3** was synthesized by modifying a previous method.[10] 2-Amino-3,5-dibromopyrazine (3000.0 mg, 12.0 mmol, 1.0 equiv.), phenylboronic acid (5800.0 mg, 48.0 mmol, 4.0 equiv.) and bis(benzonitrile)dichloro palladium (1700.0 mg, 2.4 mmol, 0.2 equiv.) were dissolved in EtOH. Next, 1N $Na_2CO_3$ (48 mL, 4.0 equiv.) was added into the reaction mixture, which was further heated to reflux overnight under Ar. After cooling down to room temperature, the mixture was filtered, and EtOH was removed with rotovap. The residue was acidified to pH 4~5 with HCl (1 N) and washed three times with EtOAc. The aqueous layer was next alkalized to pH ~11 with NaOH (1 N) and extracted three times with EtOAc. The organic layer was combined, washed with brine and $H_2O$ three times, dried with anhydrous $Na_2SO_4$, filtered, and concentrated *in vacuo* to give compound **3** (2075.0 mg, 70%).

**Diphenylterazine (DTZ):** DTZ was synthesized by modifying a previous method.[10] 6 N HCl (5.4 mL, 50.0 equiv.) was added to 5 mL of 1,4-dioxane in a 60-mL Ace pressure tube (Sigma-Aldrich #Z568767) containing compound **3** (150.0 mg, 0.61 mmol, 1.0 equiv.) and 1,1-diethoxy-3-phenylacetone (534.0 mg, 2.4 mmol, 4.0 equiv.). The tube was sealed, and the mixture was maintained at 120 °C with stirring overnight. Next, the mixture was cooled down to room temperature before the solvent was removed under reduced pressure. The crude was re-dissolved in 15 mL MeOH, and next, purified with the Waters preparative RPLC-MS (acetonitrile/water =30:70 to 98:2, 20 mL/min). Product fractions were combined and lyophilized to give DTZ (126.0 mg, 55%).

**2-Benzyl-6,8-diphenylimidazo[1,2-a]-pyrazin-3-yl prop-2-yn-1-yl succinate (Compound 4):** Compound **2** (83.0 mg, 0.53 mmol, 2.0 equiv.) and DCC (82.0 mg, 0.4 mmol, 1.5 equiv.) were placed in an oven-dried two-neck round-bottom flask purged with Ar three times. Anhydrous DCM (10 mL) mixed with TEA (80.0 µL, 0.53 mmol, 2.0 equiv.) and two drops of anhydrous DMF was injected into the reaction system with stirring at room temperature over 15 min. Subsequently, DTZ (101.7 mg, 0.27 mmol, 1.0 equiv.) was quickly added into the reaction system through one neck of the flask. The system was flushed with Ar and the reaction mixture was stirred for additional 20~30 min. The progress of the reaction was monitored with TLC (hexane/EtOAc = 3:1). After completion of the reaction, the mixture was cooled down to -20 °C to precipitate the DCU by-product. After filtration and concentration *in vacuo*, the residue was further purified with silica gel chromatography (EtOAc/hexane = 1:5, v/v) to give compound **4** (109.0 mg, 80 %). $^1$H NMR (600 MHz, $CDCl_3$) δ 8.89 – 8.84 (m, 2H), 8.24 (s, 1H), 8.16 (dd, *J* = 8.4, 1.2 Hz, 2H), 7.60 – 7.55 (m, 2H), 7.54 – 7.46 (m, 3H), 7.42 – 7.37 (m, 1H), 7.36 – 7.29 (m, 4H), 7.26 – 7.22 (m, 1H), 4.75 (d, *J* = 2.5 Hz, 2H), 4.24 (s, 2H), 2.79 – 2.74 (m, 2H), 2.73 – 2.64 (m, 2H), 2.46 (t, *J* = 2.5 Hz, 1H). $^{13}$C NMR (151 MHz, $CDCl_3$) δ 171.7, 169.0, 147.8, 138.5, 138.1, 136.8, 136.1, 135.3, 133.3, 130.4, 130.2, 129.8, 129.2, 128.8, 128.7, 128.5, 128.4, 128.3, 126.4, 126.3, 109.7, 77.2, 75.5, 75.3, 52.6, 34.3, 28.8, 28.1. ESI-MS ($C_{32}H_{25}N_3O_4$): $[M+H]^+$ calcd: 516.19, found: 516.18.

**2-(4-(((4-((2-Benzyl-6,8-diphenylimidazo[1,2-a]pyrazin-3-yl)oxy)-4-oxobutanoyl)oxy) methyl)-1H-1,2,3-triazol-1-yl)acetic acid (ETZ):** Compound **4** (83.0 mg, 0.53 mmol, 2.0 equiv.), CuI (44.0 mg, 0.23 mmol, 2.0 equiv.) and THPTA (25.0 mg, 0.23 mmol, 0.5 equiv.) were placed in a two-neck round-bottom flask purged with Ar three times. Azidoacetic acid (53.0 µL, 0.47 mmol, 4.0 equiv.) was dissolved in THF (10 mL) and injected into the reaction system. Next, ascorbic acid (41.0 mg, 0.23 mmol, 2.0 equiv.) dissolved in 2 mL dd$H_2O$ was dropwise injected into the reaction mixture and stirred overnight at room temperature. After completion of the transformation, the mixture was filtered and the product was purified with the Waters preparative RPLC-MS (acetonitrile/water = 30:70 to 98:2, v/v, 20 mL/min,). Product fractions were combined and lyophilized to give ETZ (35.0 mg, 50%). $^1$H NMR (600 MHz, methanol-$d_4$) δ 8.69 – 8.64 (m, 1H), 8.51 (s, 1H), 8.16 (dt, *J* = 6.6, 1.3 Hz, 2H), 7.92 (s, 1H), 7.57 – 7.52 (m, 3H), 7.45 (t, *J* = 7.8 Hz, 2H), 7.39 – 7.34 (m, 1H), 7.33 – 7.23 (m, 4H), 7.22 – 7.17 (m, 1H), 5.28 (s, 2H), 5.15 (s, 2H), 4.14 (s, 2H), 2.87 – 2.84 (m, 2H), 2.81 – 2.77 (m, 2H).$^{13}$C NMR (151 MHz, methanol-$d_4$) δ 178.2, 172.6, 169.7, 147.6, 142.3, 138.5, 138.1, 136.5, 135.9, 135.5, 132.9, 130.0, 129.4, 128.9, 128.7, 128.4, 128.1, 127.9, 126.1, 126.0, 125.8, 124.9, 110.1, 57.4, 50.4, 32.8, 28.3, 27.9. ESI-MS ($C_{34}H_{28}N_6O_6$): $[M+H]^+$ calcd: 617.21, found: 617.32.

**4-(2-(Dimethylamino)ethoxy)-4-oxobutanoic acid (Compound 5):** NHS (920.0 mg, 8.0 mmol, 0.4 equiv.), DMAP (260.0 mg, 2.0 mmol, 0.1 equiv.), TEA (840.0 µL, 6.0 mmol, 0.3 equiv.) and 2-dimethylaminoethanol (6.5 mL, 60.0 mmol, 3.0 equiv.) were consequently added into a solution of succinic anhydride (2000 mg, 20 mmol, 1.0 equiv) dissolved in anhydrous toluene (50 mL). The mixture was heated to reflux overnight under Ar. After cooling to room temperature, the solvent was removed *in vacuo*. The residue was dissolved in MeOH: ddH2O (1:1, v/v) and purified with the Waters preparative RPLC-MS (acetonitrile/water =3:97 to 98:2, v/v, 20 mL/min,). Product fractions were combined and lyophilized to give crude compound **5** (1580.0 mg, 42%).

**2-Benzyl-6,8-diphenylimidazo[1,2-a]pyrazin-3-yl (2-(dimethylamino)ethyl) succinate (C3-DMA-DTZ):** Compound 5 (240.0 mg, 1.3 mmol, 4.0 equiv.) and DCC (132.0 mg, 0.64 mmol, 4.0 equiv.) were placed in an oven-dried two-neck round-bottom flask purged with Ar three times. Anhydrous DCM (10 mL), DMF (1 mL), and TEA (88.0 µL, 0.64 mmol, 2.0 equiv.) were immediately added into the reaction mixture with stirring at room temperature over 30 min. Subsequently, DTZ (122.0 mg, 0.33 mmol, 1.0 equiv.) was added to the reaction mixture. The resulting mixture was stirred for additional 80 min, then filtered, concentrated, and purified with silica gel chromatography (MeOH/DCM = 1/10, v/v) to give C3-DMA-DTZ (25.1 mg, 15 %).$^1$H NMR (600 MHz, methanol-$d_4$) δ 8.68 – 8.64 (m, 2H), 8.52 (s, 1H), 8.18 – 8.14 (m, 2H), 7.58 – 7.51 (m, 3H), 7.48 – 7.44 (m, 2H), 7.40 – 7.36 (m, 1H), 7.32 – 7.26 (m, 4H), 7.23 – 7.18 (m, 1H), 4.28 (dd, *J* = 6.0, 5.2 Hz, 2H), 4.16 (s, 2H), 2.88 – 2.84 (m, 2H), 2.83 – 2.78 (m, 2H), 2.66 – 2.62 (m, 2H), 2.27 (s, 6H).$^{13}$C NMR (151 MHz, methanol-$d_4$) δ 173.2, 172.5, 130.0, 129.8, 129.4, 129.3, 128.7, 128.5, 128.4, 128.3, 128.2, 128.1, 128.0, 127.9, 127.8, 127.7, 126.2, 126.1, 126.0, 125.6, 125.5, 125.2, 58.2, 56.9, 43.9, 42.7, 32.1, 29.3, 28.2. ESI-MS ($C_{33}H_{33}N_4O_4$): $[M+H]^+$ calcd: 549.25, found: 549.10.

**BREP library construction and screening.** To create genetic fusion libraries of mScarlet-I and teLuc, pcDNA3-LumiScarlet[10] and pcDNA3-teLuc-myc[11] were used as separate PCR templates. The mScarlet-I fragment was amplified from pcDNA3-LumiScarlet with the forward primer pBAD_FW_EP and one of the three reverse primers—pBAD_RV_BREP1_NNK1, pBAD_RV_BREP1_NNK2, and pBAD_RV_BREP1_3NDT (**Supplementary Table 2**). The teLuc fragment was amplified from pcDNA3-teLuc-myc with the reverse primer pBad_RV_EP and one of the three forward primers—pBAD_FW_BREP1_NNK1, pBAD_FW_BREP1_NNK2, and pBAD_FW_BREP1_3DTZ. The amplified mScarlet-I and teLuc fragments were used for three-part Gibson assembly along with pBAD/HisB predigested with Xho I and Hind III. The resultant DNA libraries were used to transform electrocompetent E. cloni 10G cells (Lucigen), which were next allowed to grow on 2×YT agar plates containing 100 µg/mL ampicillin and 0.02% (w/v) L-arabinose at 37 °C overnight. 50 µM DTZ was sprayed onto colonies, and plates were imaged in a BLI system consisting of a UVP BioSpectrum dark box, a Computar Motorized ZOOM lens (M6Z1212MP3) and a QSI 628 Cooled CCD camera. Bioluminescence images were acquired with the µManager software. The brightest colonies were selected and cultured in 500 µL 2×YT broth supplemented with 100 µg/mL ampicillin and 0.2% (w/v) L-arabinose in 96-well deep-well bacterial culture plates. After being shaken at 30 °C, 250 rpm for 24 h, cells were pelleted by centrifugation and further lysed with 500 µL of B-PER Bacterial Protein Extraction Reagents (Pierce) at 4 °C for 30 min. Cell lysates were prepared after centrifugation. 5 µL of each lysate was further diluted with 95 µL *in vitro* assay buffer (1 mM CDTA, 0.5% Tergitol NP-40, 0.05% Antifoam 204, 150 mM KCl, 100 mM MES, pH 6.0, 1 mM DTT, and 35 mM thiourea). Next, bioluminescence spectra were recorded on a BMG Labtech microplate reader with the equipped red-sensitive PMT right after injecting 100 µL of 25 µM DTZ pre-dissolved in the assay buffer mentioned above. Mutants with high BRET efficiency and brightness were selected.

**BRIC library construction and screening.** The CaM-M13 fragment was PCR-amplified from pcDNA3.1-Orange_CaMBI_110 with oligos pBad_FW2_CaM and pBad_RV2_M13. Next, pBAD-BREP was amplified with either pBad_FW1_mScarlet and pBad_RV1_teLuc(133), or pBad_FW3_teLuc(134) and pBad_RV3_teLuc(168) to generate two fragments. The three fragments, along with pBAD/HisB predigested with Xho I and Hind III, were assembled in a four-part Gibson assembly reaction. The product was used to transform *E. coli* DH10B cells (Thermo Fisher). Cells were allowed to grow on 2×YT agar plates supplemented with 100 µg/mL ampicillin and 0.02% (w/v) L-arabinose at 37 °C overnight. Single colonies were selected for growth in 2×YT liquid culture. Plasmids were recovered and Sanger sequencing confirmed the successful creation of pBad-BRIC0.1. Next, Taq DNA polymerase (New England Biolabs) in the presence of $MnCl_2$ (0.05 mM) or GeneMorph II Random Mutagenesis Kit (Agilent Technologies) was used for error-prone PCR (EP-PCR)-based random mutagenesis. Oligos pBAD_FW_EP and pBAD_RV_EP were used for these reactions. PCR products were inserted into pBAD/HisB between Xho I and Hind III via Gibson assembly. The DNA libraries were used to transform *E. coli* DH10B cells, which were cultured on 2×YT agar plates supplemented with 100 µg/mL ampicillin and 0.02% (w/v) L-arabinose at 37 °C overnight. Red colonies were selected and used to inoculate cultures in 96-well deep-well bacterial culture plates. Each well was filled with 1 mL of 2×YT broth supplemented with 100 µg/mL ampicillin and 0.2% (w/v) L-arabinose. Cells were grown at 30 °C, 250 rpm for 48 h, pelleted by centrifugation, and lysed with 300 µL of B-PER at 4 °C for 30 min. After centrifugation, cell lysates were prepared. 5 µL of the supernatant was diluted with 185 µL of a $Ca^{2+}$-free buffer (30 mM MOPS, 100 mM KCl, 10 mM EGTA, pH 7.2) and a $Ca^{2+}$-containing buffer (30 mM MOPS, 100 mM KCl, 10 mM CaEGTA, pH 7.2) expected to provide 39 µM free $Ca^{2+}$. Next, DTZ was dissolved in the *in vitro* assay buffer described above to the concentration of 500 µM, and 10 µL of the DTZ solution was injected into each well via a reagent injector in a BMG Labtech CLARIOstar Plus microplate reader. The final concentration of DTZ was thus 25 μM. The bioluminescence spectrum and intensity of each well were measured from 400 to 700 nm with 5-nm intervals. Mutants with high brightness and $Ca^{2+}$ responsiveness were selected.

**BREP expression, purification, and emission spectrum recording.** BREP was expressed and purified following a previous procedure.[11] The purified protein was diluted with the *in vitro* assay buffer to a final concentration of 10 nM. 100 μL of the diluted protein was added into each well of a 96-well plate. 100 μL of 25 μM DTZ pre-dissolved in the *in vitro* assay buffer was injected into the well via a reagent injector in a BMG Labtech CLARIOstar Plus microplate reader. The mixture was shaken for 2 s before the bioluminescence spectrum was recorded using the equipped red-sensitive PMT. The instrument was set to scan from 400 to 700 nm with 5-nm intervals. Three technical repeats were performed to derive the average spectrum.

**BRIC and OCaMB110 expression and purification.** pBAD-BRIC was used to transform *E. coli* DH10B cells, which were next cultured on 2×YT agar plates supplemented with ampicillin (100 μg/ml). A single colony was selected and cultured in 5 mL 2×YT broth supplemented with 100 μg/mL ampicillin. The culture was shaken at 250 rpm and 37 °C overnight. The culture was next diluted with 500 mL of 2×YT medium containing 100 μg/mL ampicillin. After shaking incubation at 37 °C for 2 h, protein expression was induced by adding L-arabinose (0.2%, w/v), and the culture was maintained at 250 rpm and 16 °C for 96 h. Cells were pelleted by centrifugation and resuspended in 10 mL of 1× phosphate-buffered saline (PBS, pH 7.4) supplemented with a cOmplet Mini EDTA-free Protease Inhibitor tablet (Roche). Cells were lysed by sonication, and the lysate was clarified by centrifugation at 15,000× g for 30 min at 4 °C. The $His_6$-tagged protein was enriched with and then eluted from Ni-NTA agarose beads (Genesee Scientific). Finally, the protein was subjected to a size-exclusion HiLoad 16/600 Superdex 200 pg column (Cytiva) and eluted with an aqueous buffer containing 150 mM NaCl and 30 mM Tris HCl, pH 7.4.

A pBAD-OCaMB110 plasmid was created by amplifying the indicator gene from pcDNA3.1-Orange_CaMBI_110, but it turned to be challenging to prepare high-purity OCaMB110 protein due to proteolysis. To address the issue, an additional Strep-tag II sequence was appended to the C-terminus of the reading frame, resulting in a pBAD-OCaMB110-Step plasmid. Protein expression was performed with the identical procedure described above. To purify the protein, the N-terminal $His_6$-tagged protein in cell lysates was first enriched with Ni-NTA agarose beads, and next, the eluate was applied to a Strep-Tactin Superflow high-capacity column (IBA Lifesciences). Finally, the eluate from the Strep-Tactin column was subjected to size-exclusion chromatography. A similar pBAD-BRIC-Step plasmid was constructed in parallel. The spectral properties of the BRIC protein prepared from pBAD-BRIC-Step with dual $His_6$ and Strep tags were identical to the protein prepared from pBAD-BRIC only with the N-terminal $His_6$ tag.

**Spectroscopic characterization of BRIC and OCaMB110.** Protein concentrations were determined with the Pierce 660 nm Protein assay using bovine serum albumin (BSA) standards. The $Ca^{2+}$-containing and $Ca^{2+}$-free buffers mentioned above were used to record the emission spectra for $Ca^{2+}$-free and $Ca^{2+}$-saturated states. Final protein and substrate concentrations were 50 nM and 25 μM, respectively. The $Ca^{2+}$ affinity was determined as previously described using a series of buffers made from mixing the $Ca^{2+}$-containing and $Ca^{2+}$-free buffers.[24,40] The bioluminescence intensity at 595 nm recorded on a BMG Labtech CLARIOstar Plus microplate reader was plotted against the expected free $Ca^{2+}$ concentration in each buffer. Three technical repeats were performed, and the data was fit with a one-site binding model in GraphPad Prism8.

To determine pH sensitivity, the $Ca^{2+}$-containing and $Ca^{2+}$-free buffers were adjusted with HCl (12 M and 1M) or KOH (4 M) to gain buffers with pH values ranging from 5.5 to 8.0. Bioluminescence intensity ratios at 595 nm in the presence and absence of $Ca^{2+}$ were plotted against the pH values.

**Evaluation of luciferase substrates for brain delivery in mice.** pBad-BREP was amplified with oligos pcDNA3_FW_HindIII and pcDNA3_RV_XhoI via PCRs. The fragment was inserted into pcDNA3 to afford pcDNA3-BREP. pcDNA3-BREP was next used to transfect HEK 293T cells using a described procedure.[11] Cells were collected 20 h post-transfection and resuspended in 1×PBS (pH 7.2). 3 μL of cells (~7,000 cells) were delivered into each side of the hippocampus (AP -1.7, ML ±1.2 and DV -1.5)[41] of 8-week-old, anesthetized BALB/cJ mice (The Jackson Laboratory, #000651) via intracranial stereotactic injection at a flow rate of 200 nL $min^{-1}$. When the infusion was complete, the needle was kept in the brain for 5 min before being withdrawn. The wound was then sealed with surgical adhesive.

Meanwhile, the compounds DTZ, ETZ, and C3-DMA-DTZ were dissolved in aqueous buffers supplemented with 25% (w/v) 2-hydroxypropyl-β-cyclodextrin (HP-β-CD) and 20% (v/v) PEG-400. More specifically, a 5 mL injection buffer for DTZ was made by dissolving 1.25 g HP-β-CD and 1 mL PEG-400 in ~ 3 mL normal saline; a 5 mL injection buffer for ETZ was made by dissolving 1.25 g HP-β-CD and 1 mL PEG-400 in ~ 3 mL of normal saline pre-supplemented with 5% (w/v) $NaHCO_3$; a 5 mL injection buffer for C3-DMA-DTZ was made by dissolving 1.25 g HP-β-CD and 1 mL PEG-400 in ~ 3 mL of normal saline pre-supplemented with 100 mM citric acid (pH 5.5). $NaHCO_3$ and citric acid have been used broadly for intravenous infusion and were supplemented to formulate ETZ and C3-DMA-DTZ as salts *in situ*. This new buffer system was modified from the previously used recipe[10,14] with remarkably reduced viscosity, allowing more consistent intravenous injection.

100 μL of DTZ (2.5 mM), ETZ (6.8 mM), or C3-DMA-DTZ (4.5 mM) in their corresponding *in vitro* injection buffers was delivered to mice via tail vein right after intracranial cell injection. Bioluminescence images were acquired with a BLI system consisting of a UVP BioSpectrum dark box, a Computar Motorized ZOOM lens (M6Z1212MP3) and a QSI 628 Cooled CCD camera. The settings were: camera binning 4×4; high gain; camera sensor temperature -15 °C; 10 s exposure time and each acquisition every 60 s. The lens was controlled using the UVP VisionWorksLS software with aperture set to 100% open, zoom set to 0%, and focus set to 0%. Anesthetized mice were placed 21 cm away from the front of the lens with no emission filter used. To minimize biological variables, each mouse was sequentially tested with three substrates, and the next substrate was administered after the signal of the previous substrate faded out. A total of three mice were used, and the substrate injection order was rotated for each mouse to control the bias. Images were acquired with the μManager software and processed in the Fiji version of ImageJ 2.1. Image stacks were first subtracted for background by setting the rolling ball radius to 100 pixels. Next, the region of interest (ROI) was selected based on bioluminescence from the mouse brain, and the intensity value integrated over the ROI was extracted for further analysis. Data were plotted and statistical analysis was performed in GraphPad Prism8. After the software-based background subtraction, the images were left with residual background, so the ROI was moved away from the mouse brain region to evaluate residual background, which was further used to subtract signals for calculating the integrated bioluminescence intensity (area under the curve).

**Characterization and comparison of BRIC and OCaMBI110 in HeLa cells.** pBad-BRIC was used for PCRs with oligos pcDNA3_FW_HindIII and pcDNA3_RV_XhoI. The fragment was inserted into pcDNA3 to afford pcDNA3-BRIC. HeLa cells were transfected with 3 μg of the plasmid pcDNA3-BRIC or pcDNA3.1-Orange_CaMBI_110 using a described procedure [11]. Cells were allowed to grow at 37 °C in a 5% $CO_2$ incubator for 18 h. Cells were rinsed twice with 1× PBS and placed in 1×PBS 15 min before imaging. Images were acquired on an inverted Leica DMi8 microscope equipped with a Photometrics Prime 95B Scientific CMOS camera. 40 μM DTZ or 40 μM FRZ was supplemented for bioluminescence. Bioluminescence imaging settings were: 40× oil immersion objective lens (NA 1.2), no filter cube, 2×2 camera binning, 1 s exposure with 0 s interval, camera sensor temperature -20 °C; 12-bit and high sensitivity mode. Histamine was dissolved in 1×PBS and added during time-lapse imaging to a final concentration of 100 μM. Image stacks were processed as described in the previous section, except that ROIs were selected for individual cells and the mean of intensity values over the ROI was extracted for further analysis. Moreover, the baselines caused by substrate decay were fitted to a monoexponential model: $Y=(Y_0\text{-Plateau})*\exp(-K*X)+\text{Plateau}$. Data were plotted and statistical analysis was performed in GraphPad Prism8.

**Preparation of Adeno-Associated Viruses (AAVs).** Indicator genes were amplified from their corresponding pcDNA3/3.1 plasmids and inserted into a pAAV-hSyn vector to generate pAAV-hSyn-BRIC, pAAV-hSyn-BREP, and pAAV-hSyn-OCaMBI110. Next, individual transfer plasmids, along with pAdDeltaF6 and pAAV2/9n, were used to transfect HEK 293T cells to pack viruses. A protocol by Rego et al.[42] was followed for viral packing and purification. Viral titers were determined with quantitative PCR (qPCR) based on SYBR green by following a protocol from Addgene. Typical AAV titers were $5\times10^{14}$ - $1\times10^{15}$ GC/mL. AAVs were aliquoted and stored at -80 °C for long-term stability.

**Characterization and comparison of BRIC and OCaMBI110 in primary mouse neurons.** Freshly extracted embryonic day 18 primary mouse brain tissue was collected for neuron dissociation. The neurons were plated on poly-D-lysine coated 35 mm glass-bottom dishes with 2 mL NbActiv4 medium (BrainBits) at 37 °C and 5% $CO_2$. Half of the medium was changed to fresh NbActiv4 every 2 days. Neurons were transduced with AAVs (AAV-hSyn-BRIC, AAV-hSyn-BREP, or AAV-hSyn-OCaMBI110) at the 5th day post plating. 3 μL of each virus at ~$5\times10^{14}$ GC/mL and 1 μL of 1 M HEPES (pH 7.4) were added to each 35 mm culture dish (with cells at ~ 60% confluency). Half of the medium was replaced with fresh NbActiv 24 h later and then every 2 days. Neurons were evaluated on the fourth or fifth day after AAV transduction. Because extensive fluorescent aggregations were observed in OCaMBI110-expressing neurons on day 5, neurons on day 4 after AAV transduction were used for quantitative comparison. Growth medium was replaced with 0.8 mL of the luminescence imaging buffer[24] (0.49 mM $MgCl_2$, 2 mM $CaCl_2$, 0.4 mM $MgSO_4$, 0.44 mM $KH_2PO_4$, 5.3 mM KCl, 4.2 mM $NaHCO_3$, 0.34 mM $Na_2HPO_4$, 138 mM NaCl, 10 mM HEPES pH 7.2, 15 mM D-glucose, and 0.1 mM sodium pyruvate) supplemented with 100 μM of DTZ or FRZ. Images were acquired on an inverted Leica DMi8 microscope equipped with a Photometrics Prime 95B Scientific CMOS camera. During time-lapse imaging, 0.2 mL of the high $K^+$ stimulation buffer[24] (0.49 mM $MgCl_2$, 2 mM $CaCl_2$, 0.4 mM $MgSO_4$, 0.44 mM $KH_2PO_4$, 143.2 mM KCl, 4.2 mM $NaHCO_3$, 0.34 mM $Na_2HPO_4$, 10 mM HEPES pH 7.2, 15 mM D-glucose, and 0.1 mM sodium pyruvate) supplemented with 100 μM of DTZ or FRZ was added to depolarize cells. Instrumental settings and data analysis were identical to those described for the HeLa cell experiment, except that the exposure time was 2 s.

**Comparison of BRIC and OCaMBI110 brightness in the hippocampus in mice.** 500 nL AAV (~1× $10^{15}$ GC/mL) was delivered to each side of the hippocampus of 8-week- old BALB/cJ mice via intracranial stereotactic injection at a flow rate of 100 nL $min^{-1}$ using the coordinate described above. 19 days after viral injection, mice were utilized for brightness comparison. 100 µL ETZ (6.8 mM) in the *in vivo* injection buffer was intravenously delivered into the anesthetized BRIC-expressing mouse. Subsequently, BLI was performed with a UVP BioSpectrum dark box, a Computar Motorized ZOOM lens (M6Z1212MP3), and an Andor iXon Life 888 EMCCD camera. The camera was first set to the "Photon Counting" mode using OptAcquire pre-settings, and the gain was next adjusted down to 500. Other parameters were: camera binning 2×2; camera temperature -70 °C; 500 ms exposure time with acquisitions every 5 s. The lens was controlled using the UVP VisionWorksLS software with aperture set to 100% open, zoom set to 0%, and focus set to 0%. Anesthetized mice were placed 21 cm away from the front of the lens with no emission filter used. FRZ and FFz were dissolved in the *in vivo* injection buffer (the same recipe described above for dissolving DTZ). 100 µL of FRZ (3.0 mM) or FFz (6.0 mM) were delivered into each of the anesthetized OCaMBI11-expressing mice. Imaging conditions were unchanged. Data analysis (including residual background analysis) was identical to those described above for comparing the brain delivery of the substrates.

**BRIC responses in the acute hippocampal slices.** Acute brain slices were prepared 3 weeks post viral delivery, which is described in the previous section. Freshly extracted mouse brains were pre-cooled and sliced to be 350 µm thickness in an ice-cold ACSF buffer (2.5 mM KCl, 119 mM NaCl, 1.3 mM $MgSO_4$, 26 mM $NaHCO_3$, 1 mM $NaH_2PO_4$, 2 mM $CaCl_2$, and 10 mM glucose, 95% $O_2$/5% $CO_2$). Next, brain slices were recovered in ACSF at 37 °C for 30 min and next placed in 2.0 mL of the luminescence imaging buffer (described above for neuron experiments) supplemented with 100 µM of DTZ. Images were acquired on a Scientifica SliceScope Pro 1000 equipped with a Photometrics Prime 95B Scientific CMOS camera. Bioluminescence imaging settings were: 4× objective lens (NA 0.1), no filter cube, 2×2 camera binning, 2 s exposure with 0 s interval, camera sensor temperature -15 °C; 12-bit high sensitivity mode. During time-lapse imaging, 0.5 mL of the high-$K^+$ stimulation buffer (described above for neuron experiments) supplemented with 100 µM of DTZ was slowly added using a perfusion pump at a rate of 11.9 µL $s^{-1}$. This ~ 42 s process minimized the motion of the brain slice. BREP-expressing slices were prepared and imaged using the same procedures. Image processing and data analysis were identical to those described for the HeLa cell experiment.

***In vivo* imaging of brain activities in awake mice.** 500 nL of the virus was at a flow rate of 100 nL $min^{-1}$ delivered to each side of the basolateral amygdala (BLA) of 8-week-old BALB/cJ mice and each side of the hippocampus of 8-week-old C57BL/6J mice (The Jackson Laboratory, #000664) via intracranial stereotactic injection. The coordinate for BLA was: from Bregma, AP -1.42, ML ±3, DV -5.6.[43] The same coordinate described above was used for the hippocampus. The viral titers for BREP and BRIC were ~5 × $10^{14}$ and ~1 × $10^{15}$ GC/mL, respectively. BLI was performed two to four weeks later. For $Ca^{2+}$ dynamics in the BLA included by footshock stimuli, 100 µL ETZ (6.8 mM) in the *in vivo* injection buffer was administered to an awake mouse via tail vein. The animal was next mounted on a Narishige plastic mouse head holder (SRP-AM2).

Subsequently, BLI was performed with a UVP BioSpectrum dark box, a Computar Motorized ZOOM lens (M6Z1212MP3), and an Andor iXon Life 888 EMCCD camera. Instrumental settings were identical to the descriptions for *in vivo* BRIC and OCaMBI110 brightness comparison, except that the binning was 4×4 and the exposure was 1 s. Mice were placed 27 cm away from the front of the lens. Each experiment consisted of 100 s for animal acclimation, followed by 13 footshock trials. Each trial begins with a 0.8-mA electric footshock (lasting for 1 s), and the intervals between footshock stimuli were 40 s. The electric shock was generated with an A-M Systems 2100 isolated pulse stimulator. For $Ca^{2+}$ dynamics in the hippocampus, KA was first delivered to mice via i.p. injection at a dosage of 20 mg per kg body weight. If no seizure was observed within 2 hours, a second dose KA (5 mg per kg body weight) was delivered via i.p. injection. Finally, when evident tremble was observed (indicating the successful induction of seizures), 100 µL ETZ (6.8 mM) in the *in vivo* injection buffer was administered to the mouse via tail vein. Next, BLI was performed with the head-fixed animal as described above. Data analysis was identical to those described for the HeLa cell experiment, except that ROIs were selected for bioluminescence from the brain.

**General procedure to correct time-lapse images for baseline decays.** The Fiji version of the ImageJ 2.1 was used for image processing. Image stacks were first subtracted for background by setting the rolling ball radius to 100 pixels. Next, an average image of each stack was used to generate a binary mask, which was subsequently applied to the image stack. Thus, the information for pixels within the ROI was retained, and the intensity values for other background pixels were set to 0. This conversion was necessary so that the background signals were not amplified during the subsequent baseline decay correction. Next, the whole image stacks were subjected to "photobleaching correction". This procedure essentially fitted the image stacks with a monoexponential model: $Y=(Y_0\text{-Plateau})\times\exp(-K\times X)+\text{Plateau}$, and the intensity of each image in the stack was rescaled. This correction procedure was adequate for histamine-induced $Ca^{2+}$ dynamics in HeLa and *in vivo* imaging of footshock- and KA-stimulated mice. The imaging data for high $K^+$-induced $Ca^{2+}$ in cultured neurons and brain slices required a more complicated correction. Because high $K^+$ induced a relatively large, concerted intensity increase, the baseline was not properly identified when the whole stack was used for monoexponential fitting. Instead, the mean intensity values of each image in the stack were extracted and exported to Microsoft Excel. Data points during the expected peak responses were excluded, and monoexponential fitting was applied to the remaining data points. Finally, the offline-derived decay parameters were used to correct the whole image stacks in Fiji.

**Data and statistical analysis.** Fiji (ImageJ) was used to analyze microscopic images. Imaging background was typically subtracted by setting the rolling ball radius to 300 pixels and more detailed procedures for imaging processing are presented in the above sections. Microsoft Excel, GraphPad Prism, and Affinity Designer were used to analyze data and prepare figures for publication. Sample size and the number of replications for experiments are presented in figure legends. No statistical methods were used to pre-determine the sample size. Data are shown as mean and standard deviation (s.d.) or standard error (s.e.m), and the information is included in figure legends. *P*-values, statistical methods used to calculate *P*-values, and sample size are provided in figures or figure legends, and default settings in GraphPad Prism were used for these calculations.

**Data availability:** The plasmids pcDNA3-BREP (#172337), pcDNA3-BRIC (#172338), pAAV-hSyn-BREP (#172340), pAAV-hSyn-BRIC (#172341), and pBAD-BRIC (#172343) and their sequence information have been deposited to Addgene. Other data, materials, and methods are presented in the main text or the supplementary materials, or available from the corresponding author upon request.

#### References


39 Jiang, T. *et al.* New bioluminescent coelenterazine derivatives with various C-6 substitutions. *Org. Biomol. Chem.* **15**, 7008-7018, doi:10.1039/c7ob01554b (2017).

40 Tsien, R. & Pozzan, T. Measurement of cytosolic free $Ca^{2+}$ with quin2. *Methods Enzymol.* **172**, 230-262, doi:10.1016/s0076-6879(89)72017-6 (1989).

41 Tetteh, H., Lee, J., Lee, J., Kim, J. G. & Yang, S. Investigating Long-term Synaptic Plasticity in Interlamellar Hippocampus CA1 by Electrophysiological Field Recording. *J. Vis. Exp.*, doi:10.3791/59879 (2019).

42 Rego, M. *et al.* Improved yield of AAV2 and rAAV2-retro serotypes following sugar supplementation during the viral production phase. *bioRxiv*, DOI: 10.1101/488585, doi:10.1101/488585 (2018).

43 Unger, E. K. *et al.* Directed Evolution of a Selective and Sensitive Serotonin Sensor via Machine Learning. *Cell* **183**, 1986-2002.e1926, doi:10.1016/j.cell.2020.11.040 (2020).

#### SUPPLEMENTARY INFORMATION

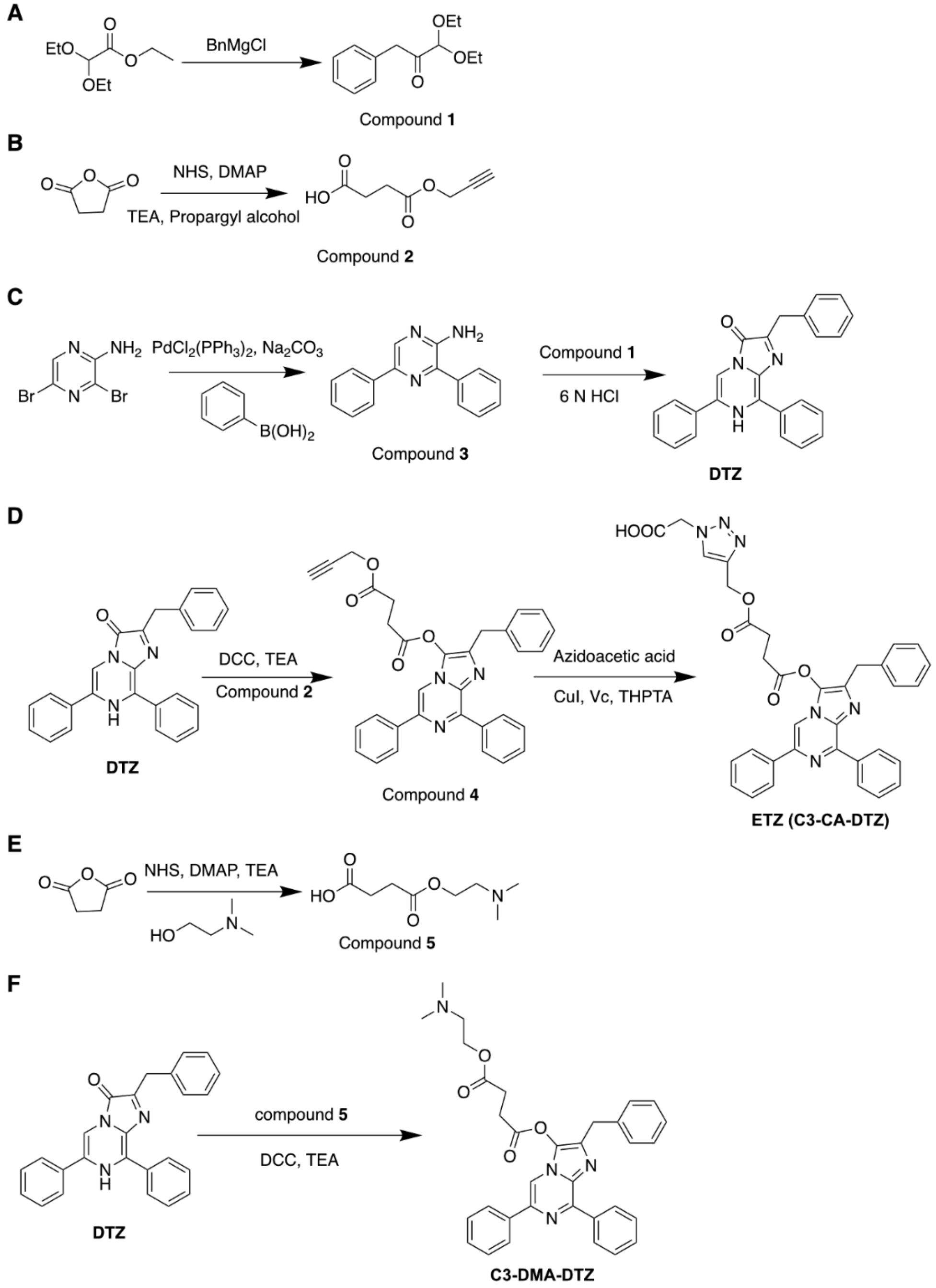


**Supplementary Fig 1. Chemical reactions for preparing ETZ and C3-DMA-DTZ.** The detailed procedures are described in the Materials and Methods section. NMR spectra for key compounds are included at the end of this document. (**A**) Anhydrous THF, -78 °C 1 h, then RT 1 h, 70%. (**B**) Anhydrous toluene, 110 °C overnight, 65%. (**C**) Step 1: EtOH, reflux overnight, 70%. Step 2: 1, 4-dioxane, 90 °C overnight, 55%. (**D**) Step 1: Anhydrous DCM, RT 30 min, 80%. Step 2: THF and ddH$_2$O (5:1), RT overnight, 50%. (**E**) Anhydrous toluene, 110 °C overnight, 42%. (**F**) Anhydrous DCM and DMF (10:1), RT 80 min, 15%.

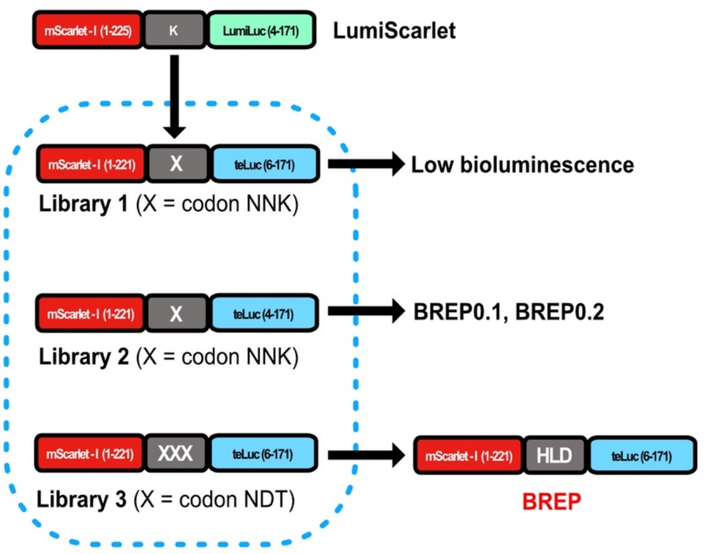


**Supplementary Fig 2. Schematic illustration of the workflow to engineer BREP.** The numbers in parentheses indicate amino acid residue numbers in the initial proteins. For codons, N refers to A, T, G, or C; K refers to G or T; D refers to A, G, or T. BREP contains a three-residue (HLD) linker and truncations of the C-terminus of mScarlet-I and the N-terminus of teLuc. Considering these truncations, mScarlet-I and teLuc in BREP are positioned closer by four residues than mScarlet-I and LumiLuc in LumiScarlet.

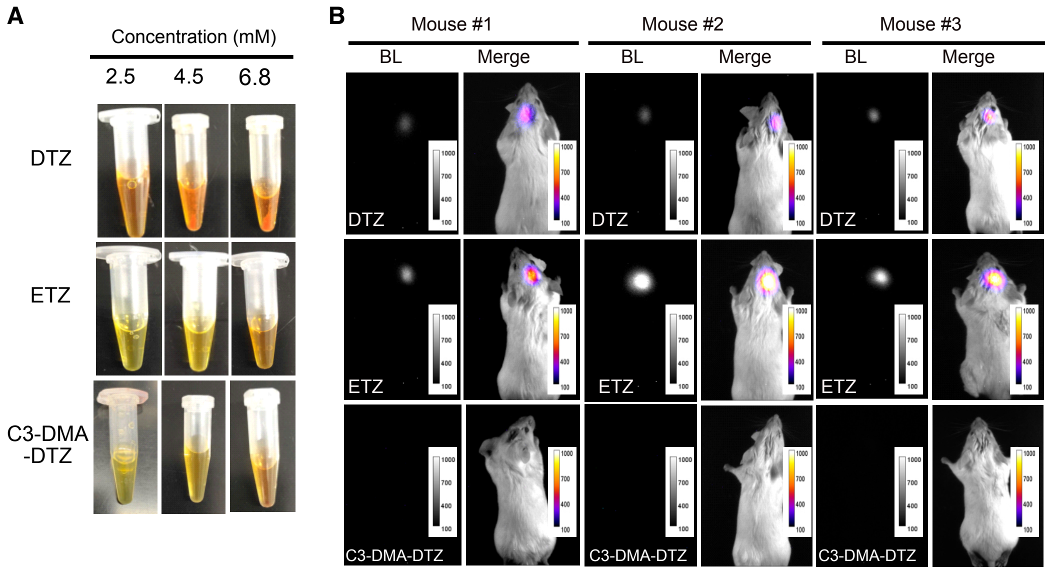


**Supplementary Fig 3. Comparison of the synthetic luciferase substrates (DTZ, ETZ and C3-DMA-DTZ) for brain imaging.** (**A**) Solubility tests of the substrates in the intravenous injection buffers, showing the approximate solubility of DTZ, ETZ and C3-DMA-DTZ to be 2.5, 6.8, or 4.5 mM, respectively. (**B**) BLI of live mice with BREP-expressing HEK 293T cells stereotactically injected into the hippocampus. The substrates were administered via tail vein at their respective, saturation concentrations. Images with peak bioluminescence intensities from each substrate injection were shown in grayscale (left) or pseudocolor overlaid on corresponding brightfield images (right). A group of the images is also presented in Fig. 1D.

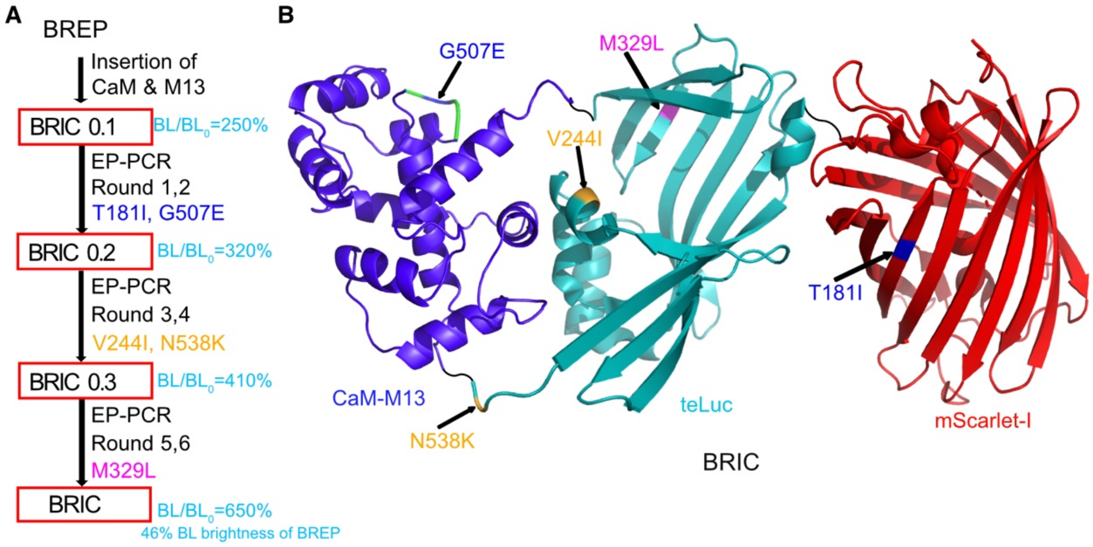


**Supplementary Fig 4. Engineering and structural illustration of BRIC.** (**A**) Flowchart to show the multistep process to develop and optimize BRIC. Calmodulin (CaM) and M13 were inserted between residues 133 and 134 of teLuc in BREP to create BRIC0.1. Next, six rounds of error-prone (EP)-PCRs were performed to enhance brightness and $Ca^{2+}$-induced responsiveness, leading to the final BRIC variant with a 6.5-fold ($BL/BL_0$) turn-on response. $BL/BL_0$ and mutations gained during the engineering process are also presented. (**B**) Schematic illustration of the domain arrangement of BRIC. Protein Data Bank entries 2BBM, 7MJB, and 5LK4 were used to create this graph. Gained mutations are highlighted.

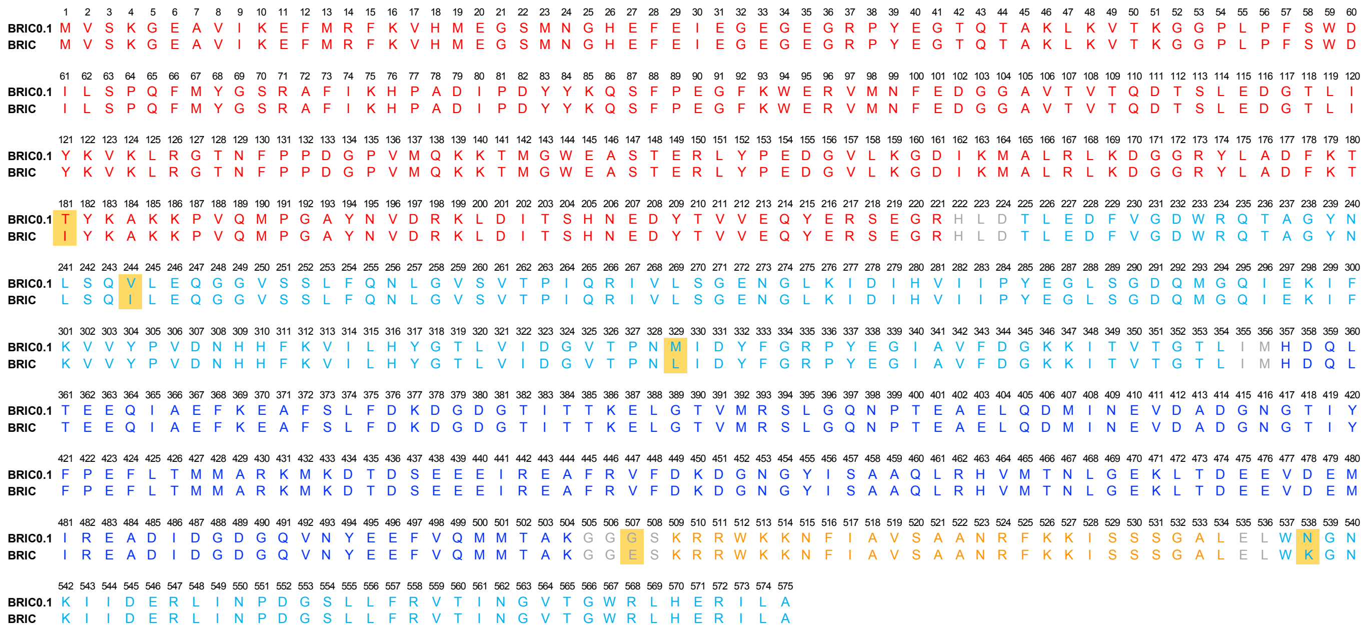


**Supplementary Fig 5. Sequence alignment of BRIC and BRIC0.1.** Sequences derived from mScarlet-I, teLuc, calmodulin, and M13 are colored in red, cyan, blue, and orange, respectively. Linker residues are colored in gray. Mutations gained during directed evolution are shaded in yellow.

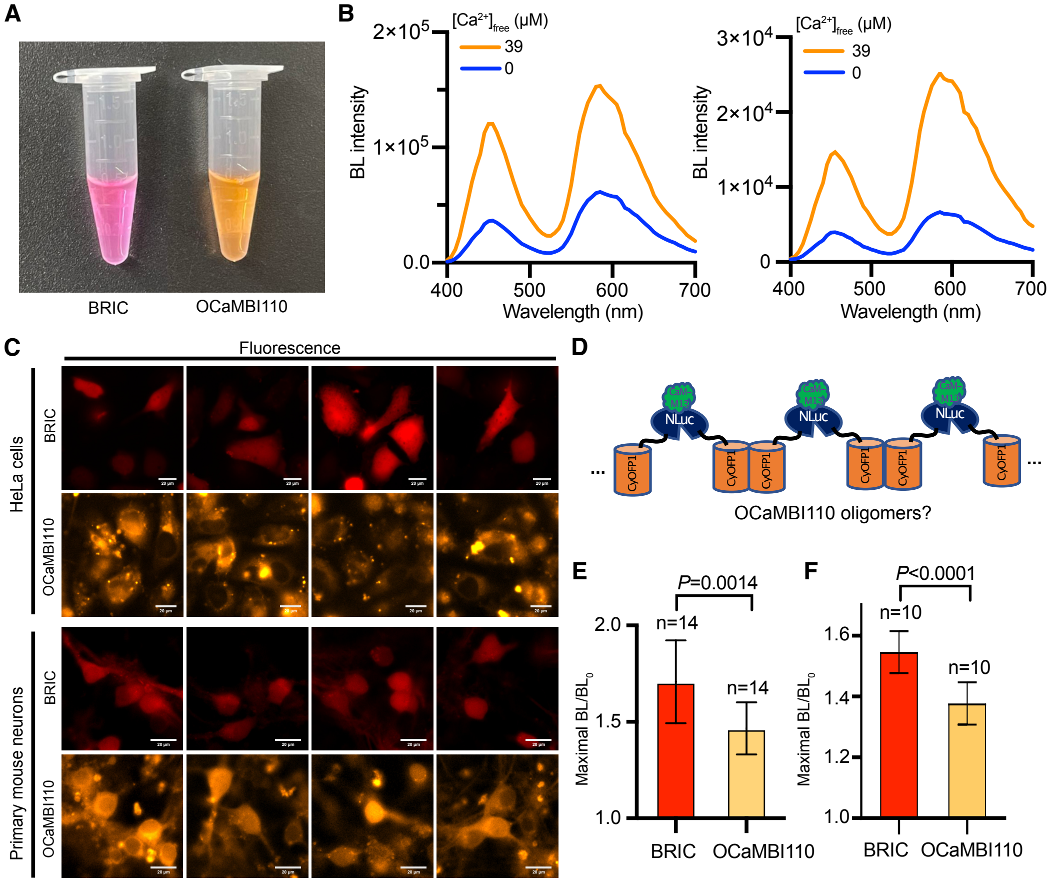


**Supplementary Fig 6. *In vitro* characterization of OCaMBI110 and comparison with BRIC in HeLa cells and primary mouse neurons.** (**A**) Photo of the two purified proteins. (**B**) Bioluminescence spectra of OCaMBI110 in the presence of FRZ and the indicated concentrations of free $Ca^{2+}$. A maximal 2.5-3.7 fold of bioluminescence increase ($BL/BL_0$) was observed using different protein preps. (**C**) Multiple views of BRIC- and OCaMBI110-expressing HeLa cells and primary mouse neurons under the fluorescence channels (Scale bar, 20 μm). Extensive fluorescent puncta were observed in cells overexpressing OCaMBI110 (*e.g.*, HeLa cells 18 h after transfection, and neurons on day 5 after AAV transduction), and these cells with fluorescent puncta showed little bioluminescence activity and were unresponsive to histamine or high $K^+$. The problem was not seen in BRIC-expressing cells. (**D**) Illustration of the possible formation of oligomers from OCaMBI110, in which CyOFP1 may form intermolecular dimers to bridge individual OCaMBI110 units. (**E**,**F**) Comparison of BRIC and OCaMBI110 for histamine-induced $Ca^{2+}$ in HeLa cells (E) and high $K^+$-induced depolarization in primary mouse neurons (F). The ratio of the maximal bioluminescence post treatment to the initial intensity is used for comparison. OCaMBI110-expressing HeLa cells were pre-selected to exclude those with fluorescent puncta. In addition, the experiment used neurons on day 4 after AAV transduction, when OCaMBI110 puncta were not obvious. Data are presented as mean±s.d., and $P$ values were derived from unpaired two-tailed $t$-tests. The BRIC data are also used for comparison with BREP in Fig. 2G.

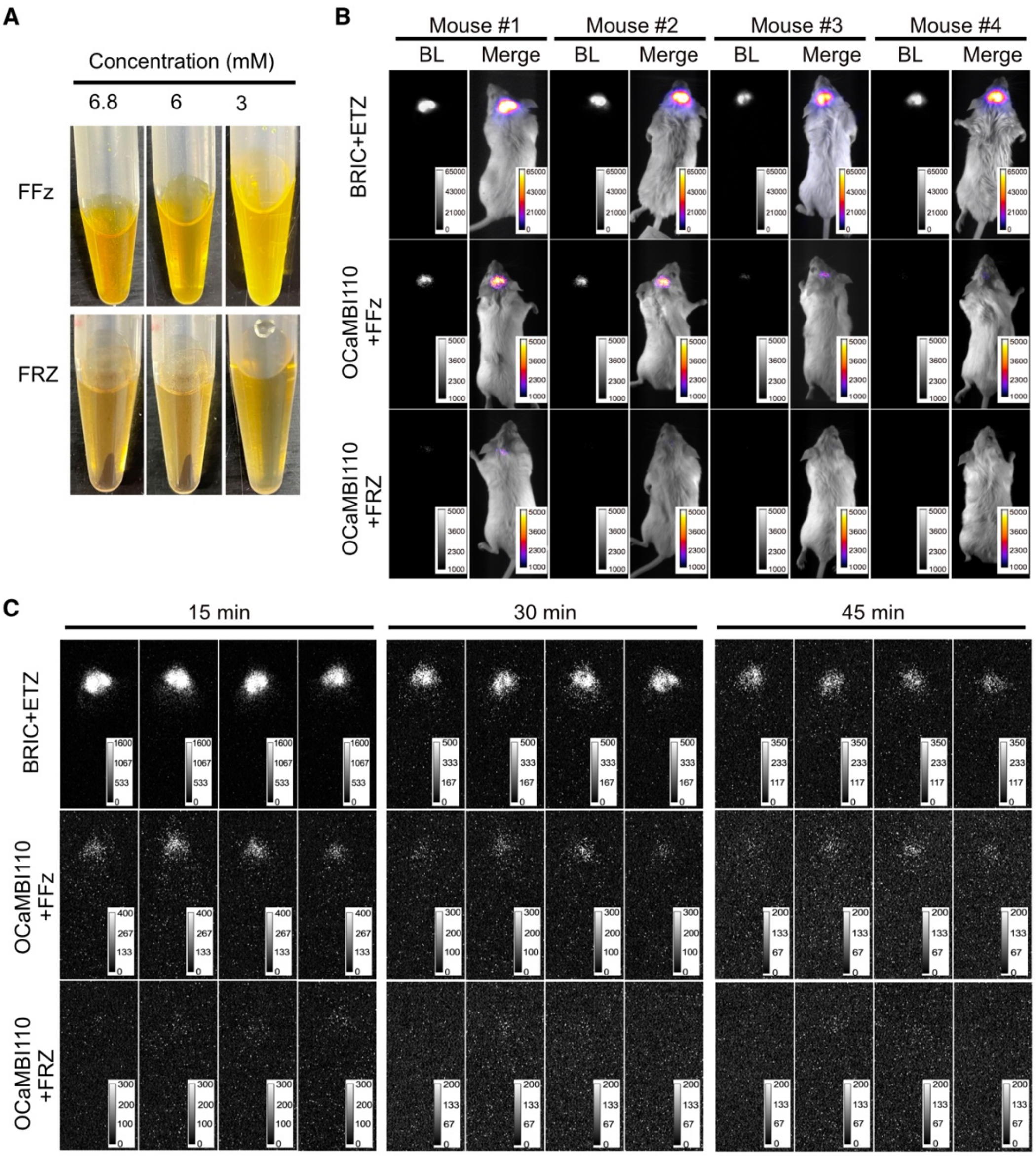


**Supplementary Fig 7. Brightness comparison of BRIC and OCaMBI110 in the hippocampus in live mice.** (**A**) Solubility tests of FFz and FRZ in the intravenous injection buffers, showing their approximate solubility to be 6 and 3 mM, respectively. (**B**) Bioluminescence images of live mice with the hippocampus transduced with BRIC or OCaMBI110 AAVs. Mice in the BRIC group were administered 6.8 mM ETZ via tail vein. Mice in the OCaMBI110 groups were administered 6 mM FFz or 3 mM FRZ via tail vein. Images at the beginning of individual imaging sessions (right after substrate injection) were shown in grayscale (left) or pseudocolor overlaid on corresponding brightfield images (right). A group of the images is also presented in Fig. 3. (**C**) Bioluminescence images in grayscale at 15, 30, and 45 min post substrate injections. In panels B and C, results from different experimental groups are presented with varying intensity ranges due to large intensity differences.

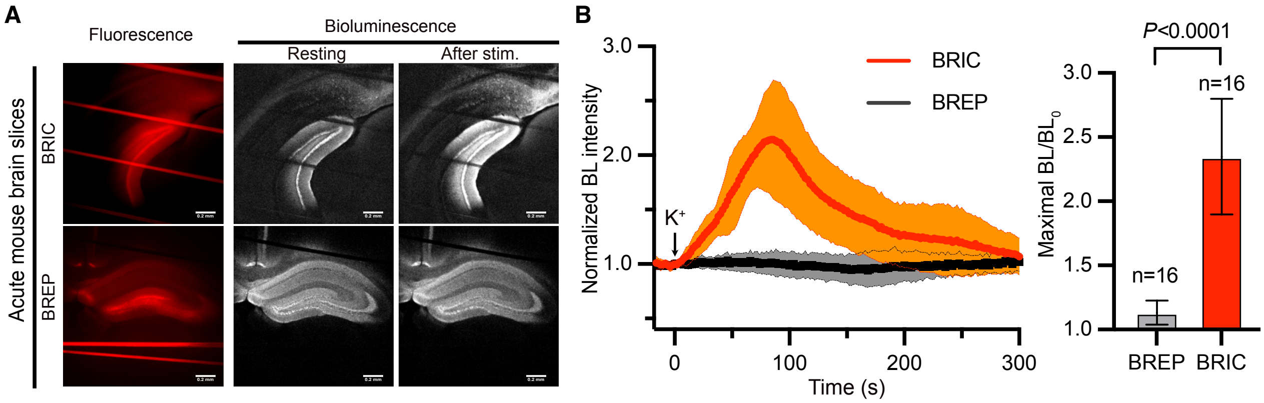


**Supplementary Fig 8. Bioluminescence imaging of high-K$^+$-induced Ca$^{2+}$ in acute mouse hippocampal slices.** (**A**) Reprehensive fluorescence and bioluminescence images of brain slices expressing BRIC or BREP (Scale bar, 0.2 mm). A peristaltic pump was used to introduce a high KCl buffer via a 42-s period. The final K$^+$ concentration was 30 mM. (**B**) Quantification of bioluminescence intensities of brain slices in response to high K$^+$. The baselines were corrected using a monoexponential decay model. Data are presented as mean±s.d., n=16 randomly chosen areas from 5 slices. *P* values were derived from unpaired two-tailed *t*-tests.

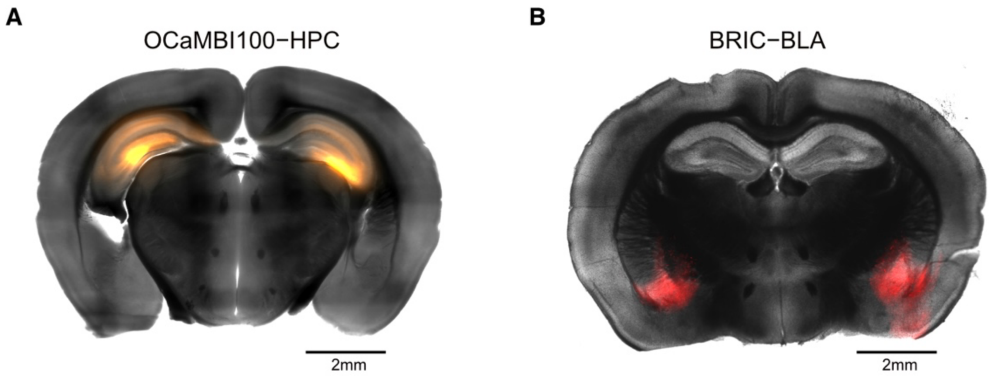


**Supplementary Fig 9. Image of acute brain slices prepared from AAV-transduced mice.** (**A**) An acute brain slice prepared from a mouse with OCaMBI110 virus injected to the hippocampus (HPC). (**B**) An acute brain slice prepared from a mouse with BRIC virus injected to the basolateral amygdala (BLA). The fluorescence channel (orange or red) is overlaid on the corresponding grayscale brightfield image. Scale bar, 2 mm.

#### Supplementary Table 1. Properties of bioluminescence $Ca^{2+}$ indicators based on NanoLuc or NanoLuc-derived luciferases.[a]

| Construct | Peak emission (nm) [b] | $BL/BL_0$ [c] | $K_d$ (nM) | Emission fraction > 600 nm | BRET Donor | BRET Acceptor | $Ca^{2+}$ sensing domain | Reference |
|---|---|---|---|---|---|---|---|---|
| GeNLs($Ca^{2+}$) | 517 | 3.9-5 | 60-520 | 0.017[d] | NLuc split at residue 66 | mNeonGreen | CaM, M13 | [19] |
| CalFluxVTN | 525 | 5[e] | 480 | 0.04[d] | NLuc | Venus | TnC | [20] |
| LUCI-GECO1 | 515 | 2.6[e] | 285 | <0.02 | NLuc | ncpGCaMP6s | CaM, RS20 | [21] |
| GLICO | 515 | 23 | 230 | <0.02 | NanoBiT | GCaMP6f | CaM, RS20 | [22] |
| ReLICO | 452 | 3.4 | $1.5\times10^6$ | 0.008 | NanoBiT | R-CEPIA1*er* | CaM, RS20 | [22] |
| Orange CaMBIs | 586 | 7-8[f] | 110-300 | 0.33[d] | NLuc split at residue 133 | CyOFP1 | CaM, M13 | [24] |
| BRIC | 595 | 6.5 | 133 | 0.54 | teLuc split at residue 133 | mScarlet-I | CaM, M13 | This work |

[a]Data for BRIC were determined in this work. Unless otherwise indicated, other data were reported in or calculated from graphs in the initial publications. [b]Wavelength for the major or the most red-shifted emission peak. [c]Intensity ratio with or without $Ca^{2+}$ at the indicated peak emission wavelength. [d]Adapted from Reference [24]. [e]Originally described as green/blue ratiometric indicators, and the response range ($R/R_0$) was reported to be 11 and 5 for CalFluxVTN and LUCI-GECO, respectively. [f]Our determined value is ~ 2.5-3.7 for OCaMBI110. The discrepancy may be caused by expression and assay conditions and the variable oligomerization states of Orange CAMBIs.

#### Supplementary Table 2. Oligonucleotides used in this work

| Oligo name | Sequence (5′→3′) |
|---|---|
| pBAD_FW_BREP1_NNK1 | CAGTACGAACGCTCCGAGGGCCGCNNKGAAGATTTCGTTGGGGAC |
| pBAD_RV_BREP1_NNK1 | GTCCCCAACGAAATCTTCMNNGCGGCCCTCGGAGCGTTCGTACTG |
| pBAD_FW_BREP1_NNK2 | CAGTACGAACGCTCCGAGGGCCGCNNKACACTCGAAGATTTCGTTGGGGAC |
| pBAD_RV_BREP1_NNK2 | GTCCCCAACGAAATCTTCGAGTGT MNNGCGGCCCTCGGAGCGTTCGTACTG |
| pBAD_FW_BREP_3NDT | CAGTACGAACGCTCCGAGGGCCGCNDTNDTNDTACACTCGAAGATTTCGTTGGGGAC |
| pBAD_RV_BREP_3NDT | GTCCCCAACGAAATCTTCGAGTGTAHNAHNAHNGCGGCCCTCGGAGCGTTCGTACTG |
| pBad_FW1_mScarlet | GACGATAAGGATCCGAGCTCGAGCATGGTGAGCAAGGGCGAG |
| pBad_RV1_teLuc(133) | CTTCTGTCAGTTGGTCATGCATAATCAGGGTCCCTGTTACAGTG |
| pBad_FW2_CaM | CACTGTAACAGGGACCCTGATTATGCATGACCAACTGACAGAAG |
| pBad_RV2_M13 | GATAATTTTGTTGCCTTTCCAGAGCTCCAGTGCCCCGGAGCTGGAGA |
| pBad_FW3_teLuc(134) | TCTCCAGCTCCGGGGCACTGGAGCTCTGGAAAGGCAACAAAATTATC |
| pBad_RV3_teLuc(168) | TCTCATCCGCCAAAACAGCCAAGCTTTTACGCCAGAATGCGTTCATG |
| pBad_FW_EP | ATGACGATAAGGATCCGAGCTCGAG |
| pBad_RV_EP | CTCATCCGCCAAAACAGCCAAGCTTTTA |
| pBad_FW_BRIC | CCGCTCGAGCATGGTGAGCAAGGGCGAGG |
| pBad_RV_strep(BRIC) | CCCAAGCTTTTATTTTTCGAACTGCGGGTGGCTCCACGCCAGAATGCGTTCATGC |
| pBad_FW_His(CaMBI) | ATAAGGATCCGAGCTCGAGCATGGTGAGCAAGGGCGAG |
| pBad_RV_strep(CaMBI) | TTTTTCGAACTGCGGGTGGCTCCACTTATAGAGTTCATCCATTC |
| pBad_RV_ext | TCATCCGCCAAAACAGCCAAGCTTTTATTTTTCGAACTGCGGGTGGCTCCAC |
| pcDNA3_FW_HindIII | TACGACTCACTATAGGGAGACCCAAGCTTGCCACCATGGTGAGCAAGGGCGAGGCAG |
| pcDNA3_RV_XhoI | TAGGGCCCTCTAGATGCATGCTCGAGTTACGCCAGAATGCGTTCATGCAGAC |
| pAAV_hSyn_FW(BRIC/BREP) | ATTCAAGCTGCTAGCAAGGATCCCGCCACCATGGTGAGCAAGGGCGAG |
| pAAV_hSyn_RV(BRIC/BREP) | TCCAGAGGTTGATTATCGATAAGCTTTTACGCCAGAATGCGTTCATGCAGAC |
| pAAV_hSyn_FW(CaMBI) | ATTCAAGCTGCTAGCAAGGATCCCGCCACCATGGTGAGCAAGGGCGAG |
| pAAV_hSyn_RV(CaMBI) | TCCAGAGGTTGATTATCGATAAGCTTTTACTTATAGAGTTCATCCATTC |

#### Captions for Movies

**Supplementary Movie 1.**
BLI of BRIC-expressing HeLa cells in response to histamine.

**Supplementary Movie 2.**
BLI of BRIC-expressing mouse neurons in response to high-$K^+$ depolarization.

**Supplementary Movie 3.**
BLI of a BRIC-expressing hippocampal brain slice in response to high-$K^+$ depolarization.

**Supplementary Movie 4.**
BLI of a live mouse with BRIC expressed in the BLA in response to 13 repeats of footshock stimulation.

**Supplementary Movie 5.**
BLI of a live mouse with BRIC expressed in the hippocampus during KA-induced seizures.

### NMR Spectra

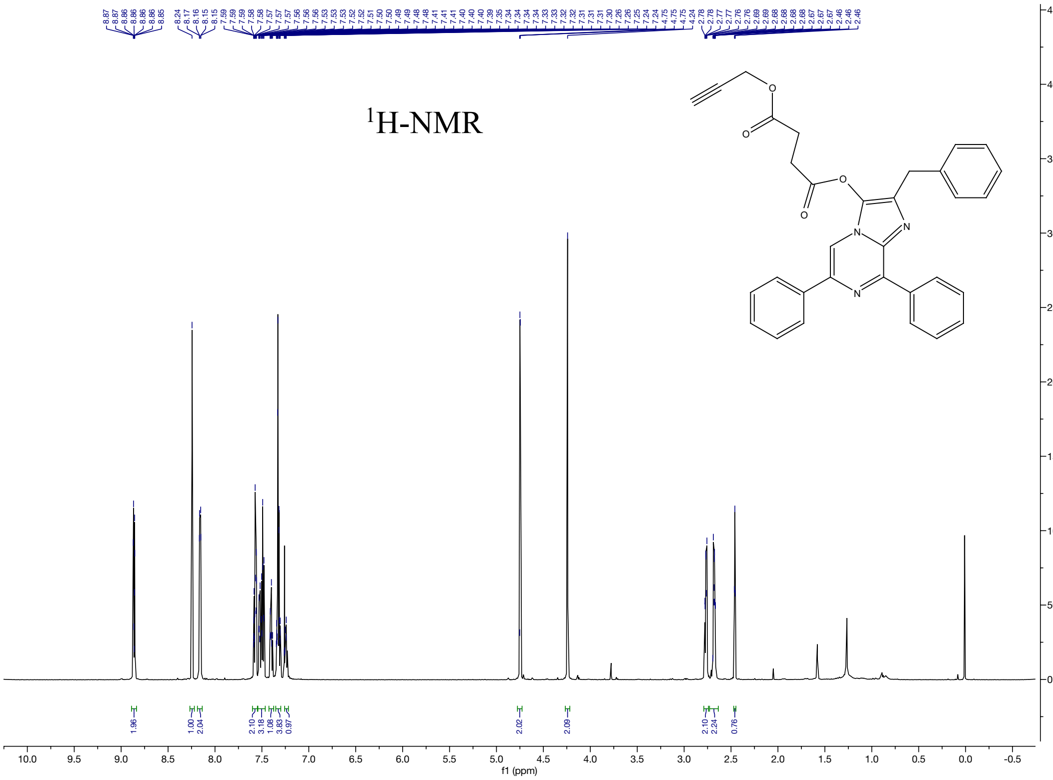


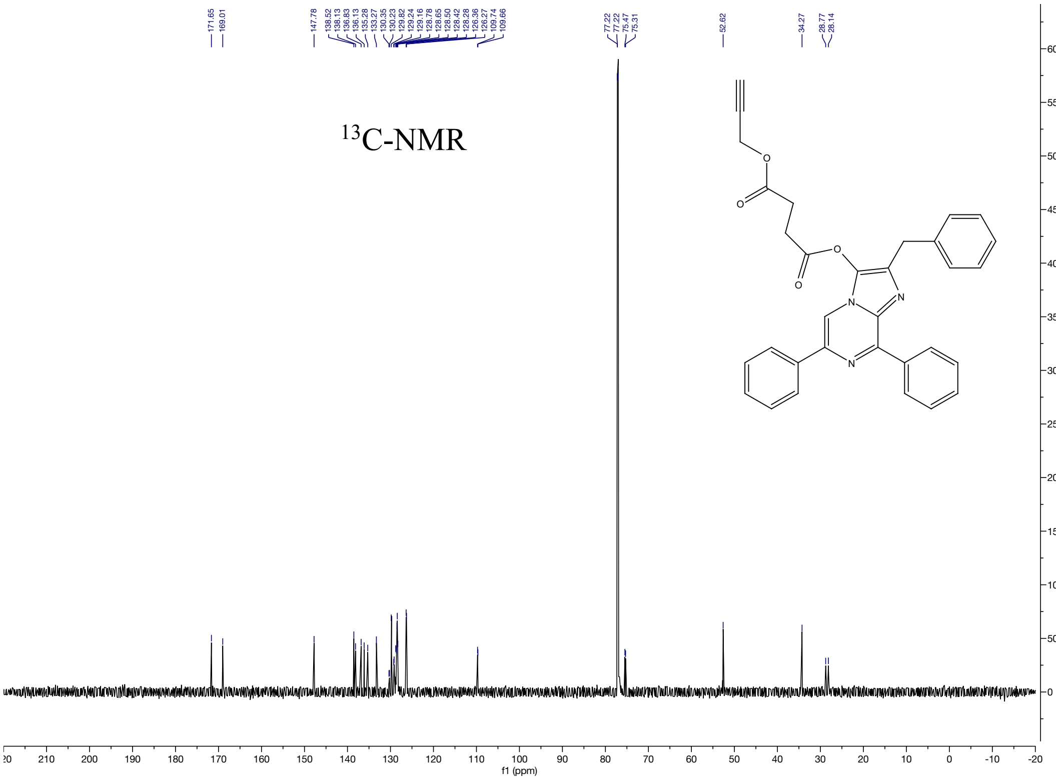

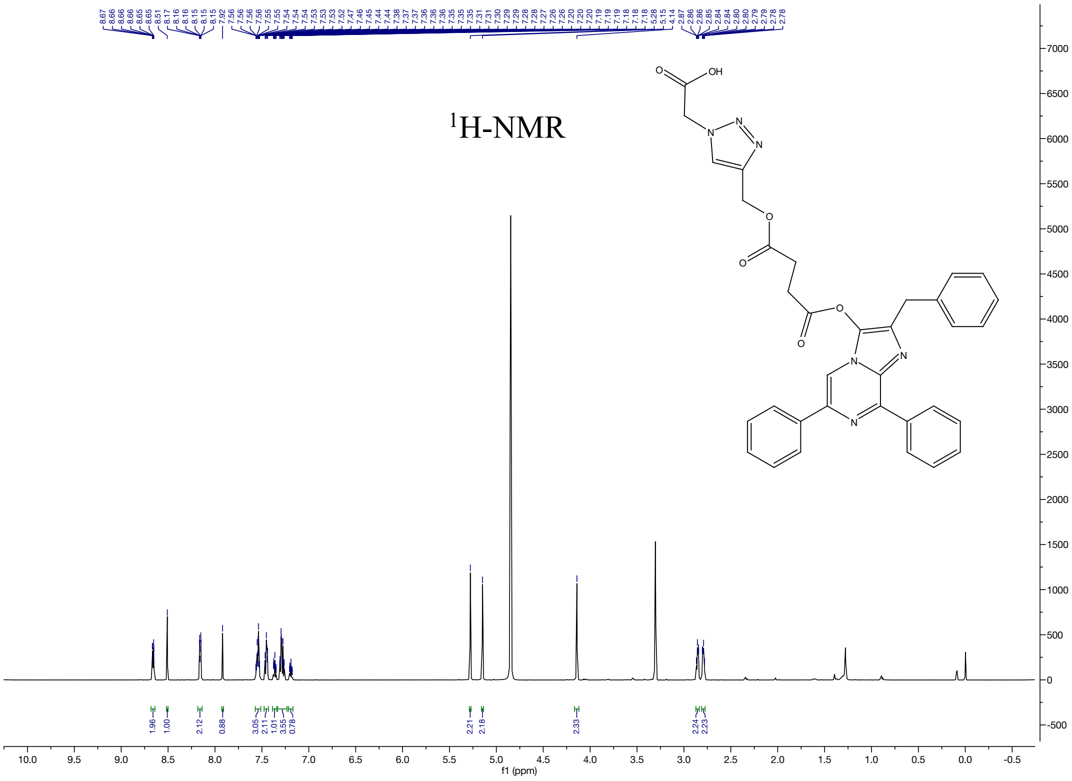
$^1$H-NMR
f1 (ppm)

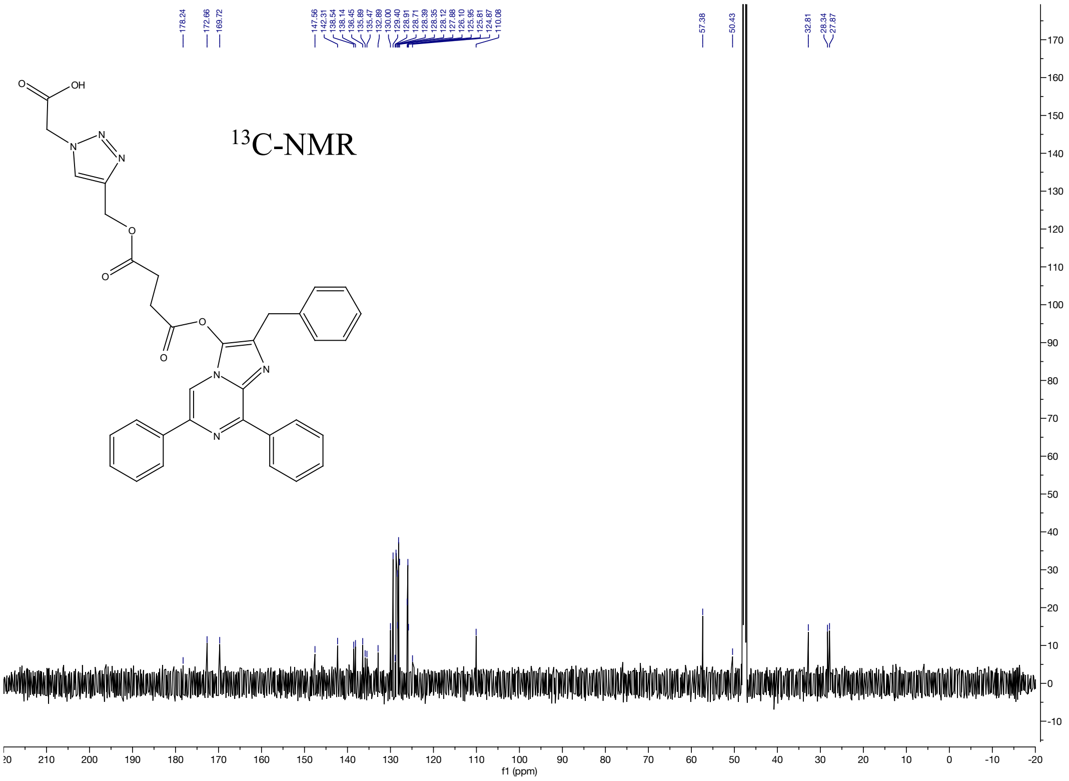
$^{13}$C-NMR
f1 (ppm)

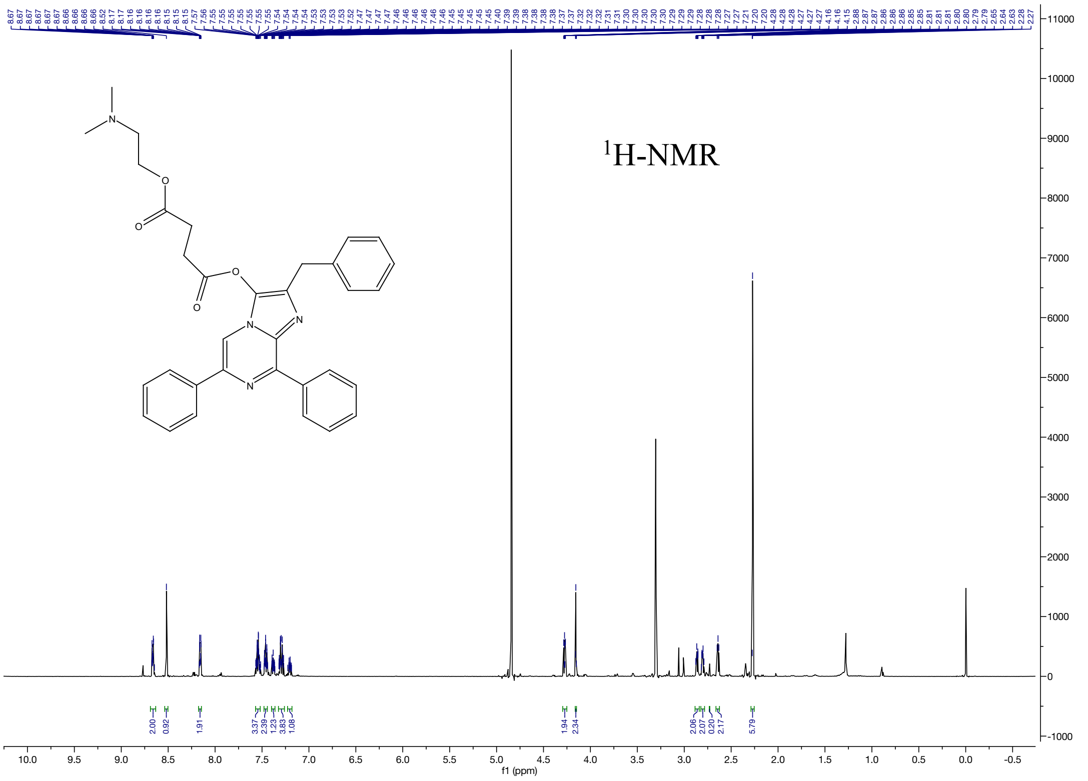

1H-NMR


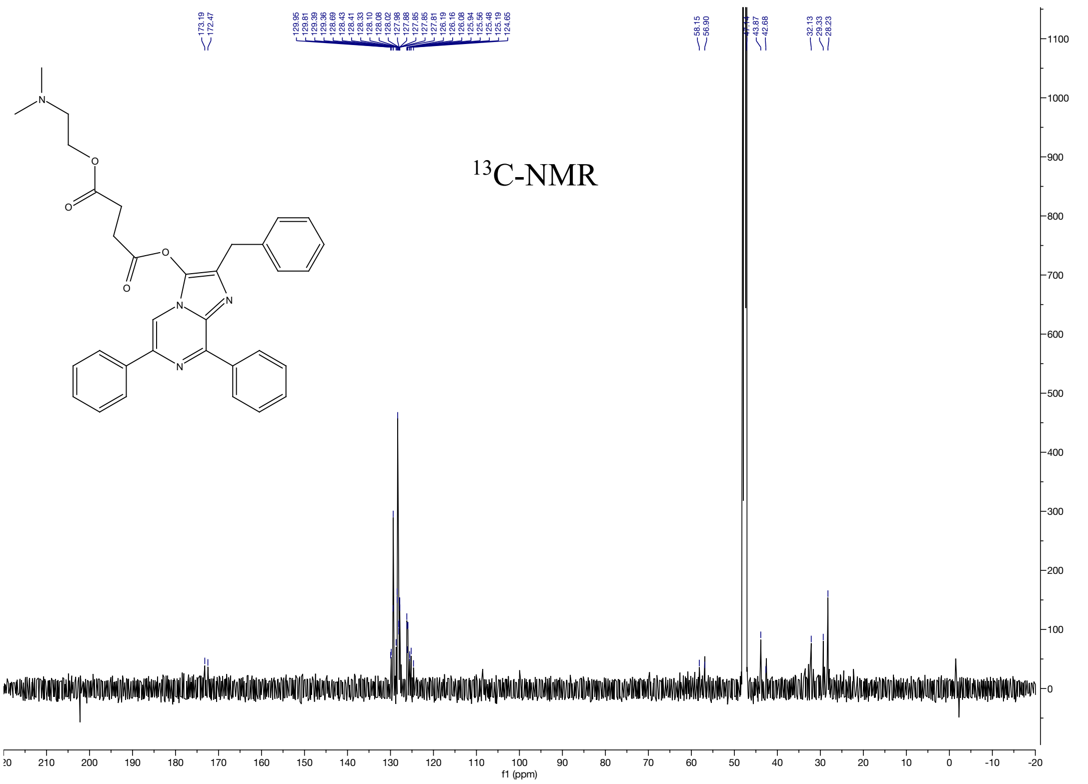

13C-NMR
